# Neural and behavioral organization of rapid eye movement sleep in zebrafish

**DOI:** 10.1101/2023.08.28.555077

**Authors:** Vikash Choudhary, Charles R. Heller, Sophie Aimon, Lílian de Sardenberg Schmid, Drew N. Robson, Jennifer M. Li

## Abstract

Sleep is ubiquitous across the animal kingdom. However, the definition for sleep varies significantly across species. In organisms such as *C. elegans, Drosophila*, and zebrafish, sleep is classically defined as periods of locomotor quiescence with an increased arousal threshold^1–6^. In mammals, sleep is further subdivided into rapid eye movement (REM) and non-rapid eye movement (NREM) sleep, but the evolutionary origin of these two sleep states remains unclear^7–12^. Using longitudinal, high-resolution imaging of naturalistic sleep-wake behavior, we find that the classical definition of larval zebrafish sleep encompasses two clearly distinct quiescent states, one with rapid eye movements (qREM) and one without rapid eye movements (qNREM). Remarkably, qREM states are maintained in congenitally blind larval zebrafish, suggesting that qREM emerged during vertebrate evolution not as a response to internally generated visual imagery (i.e., the “scanning hypothesis”)^13^, but potentially as a strategy to maintain oculomotor stimulation during rest^14^. Lastly, using brain-wide imaging in freely swimming animals, we find that, in contrast to fixed point attractor models of sleep developed in invertebrates^15^, population-level activity during zebrafish qREM unfolds along smooth committed trajectories through state-space. This work suggests a significant rethinking of the behavioral and neural organization of sleep in early vertebrates and the emergence of REM during vertebrate evolution.

## INTRODUCTION

The definition of sleep and the associated complexity of sub-states within sleep vary greatly across species. In animals such as *Drosophila melanogaster* and *Danio rerio* (zebrafish), sleep bouts are classified as prolonged locomotor quiescence (i.e., >= 1 min of immobility) and increased arousal threshold^3–6^. In mammals, sleep can be further subdivided into two major sub-states – Rapid Eye Movement (REM) / Paradoxical Sleep (PS) and Non-Rapid Eye Movement (NREM) / Slow Wave Sleep (SWS)^16^. REM sleep, with its unique behavioral signature of saccadic eye movements, was first identified in 1953^17^ and has since been the subject of debate as to its evolutionary origin, function, and underlying neural dynamics^14,18–25^.

Behavioral evidence for REM in non-mammalian species has been inconsistent. Recent studies have reported eye movements in octopuses^26^ or retinal twitches in spiders during quiescence^27^, suggesting that REM may have emerged quite early in evolution. Surprisingly, no eye movements have yet been observed during locomotor quiescence in larval or adult teleost fish^28,29^, suggesting that while some invertebrate species may have evolved a REM state independently through convergent evolution, vertebrate REM emerged later in evolution, between fish and reptiles^11^. However, no study to date has simultaneously recorded high resolution eye movements and locomotion across a 24 hr circadian cycle under naturalistic conditions in freely swimming teleost fish. Thus, the naturally occurring progression of fish sleep state(s) and their behavioral signatures remains unclear.

In parallel to the debate over the evolutionary origin of REM, there are several competing hypotheses for why eye movements during quiescence exist: the “scanning hypothesis”, which posits that saccadic eye movements are driven by internally generated visual imagery^13^, and the oculomotor / motor learning hypotheses, which posit that eye movements and body twitches during REM sleep serve to stimulate, balance, and develop motor systems during rest^14,30,31^. One way to distinguish between these hypotheses is to examine REM sleep from birth in visually deprived or blind animals. However, only a limited number of such studies have been performed across species^32,33^.

Lastly, though numerous brain regions and neuromodulators have been implicated in regulation of quiescence across species, comprehensive imaging of brain-wide neural activity at cellular resolution in freely behaving animals has been so far limited to *C. elegans*^15^. In *C. elegans*, neural activity collapses to a fixed-point attractor state during sleep, characterized by global activity suppression. A fixed-point attractor model has significant implications for the nature of sleep and wake transitions (e.g., state transitions must be stochastic), but it is unclear to what extent this model is generally true across species.

To address these questions, we used a new large-scale behavioral imaging system to simultaneously observe naturalistic behavior in freely moving animals (up to 20 fish per experiment) at high resolution (20-50 μm/pixel), across circadian timescales (e.g., 24 hours) as animals naturally alternated between periods of locomotor exploration and quiescence. Within the quiescent state, we found strong evidence for two distinct sub-states – a quiescent state with Rapid Eye Movements (qREM), and a quiescent state with No Rapid Eye Movements (qNREM). qREM is associated with behavioral immobility, an increased arousal threshold, and loss of postural control, consistent with the established definition of sleep in zebrafish^3,4,6^. Surprisingly, qREM and qNREM are coupled to distinct phases of the circadian cycle, suggesting a novel organization of sleep states that has not been previously described in other species. Finally, we found that qREM states remain intact in congenitally blind larval zebrafish, which argues against the scanning hypothesis of REM and in favor of the oculomotor hypothesis.

Using brain-wide imaging in freely swimming animals, we monitored neural activity during natural transitions between wake and qREM. We found that brain activity is broadly suppressed during qREM, with associated suppression of the noradrenergic cells in the locus coeruleus, akin to mammals^34–36^. In contrast to the previously established fixed point attractor model in *C. elegans*, neuronal subpopulations remain active during qREM and evolve along organized, committed trajectories through the state-space of population activity, raising the possibility that distinct neural dynamics emerged in the evolution of vertebrate sleep.

## RESULTS

### Identification of a novel quiescent state in larval zebrafish

Studies of larval zebrafish quiescence have primarily measured bulk locomotor movement in freely swimming animals ^3,4,6^ or eye movements in immobilized animals^28,37^, but not both simultaneously. We developed a set of new imaging systems for high resolution, high speed, longitudinal recording of behavior (**Methods: DASHER**), enabling us to simultaneously track both locomotion and eye saccades (**Fig. 1a, Supplementary Video 1**) in freely swimming larval zebrafish. We calculated average speed (mm/s) throughout the experiment and observed that fish spontaneously alternated between periods of high locomotor activity and quiescence, each often lasting on the order of tens of minutes (**Fig. 1b**). Thus, quiescent states broadly fit the definition of zebrafish sleep bouts established in previous studies (i.e., no detectable locomotor movement for >= 1 min)^3,4,6^.

**Figure 1.**
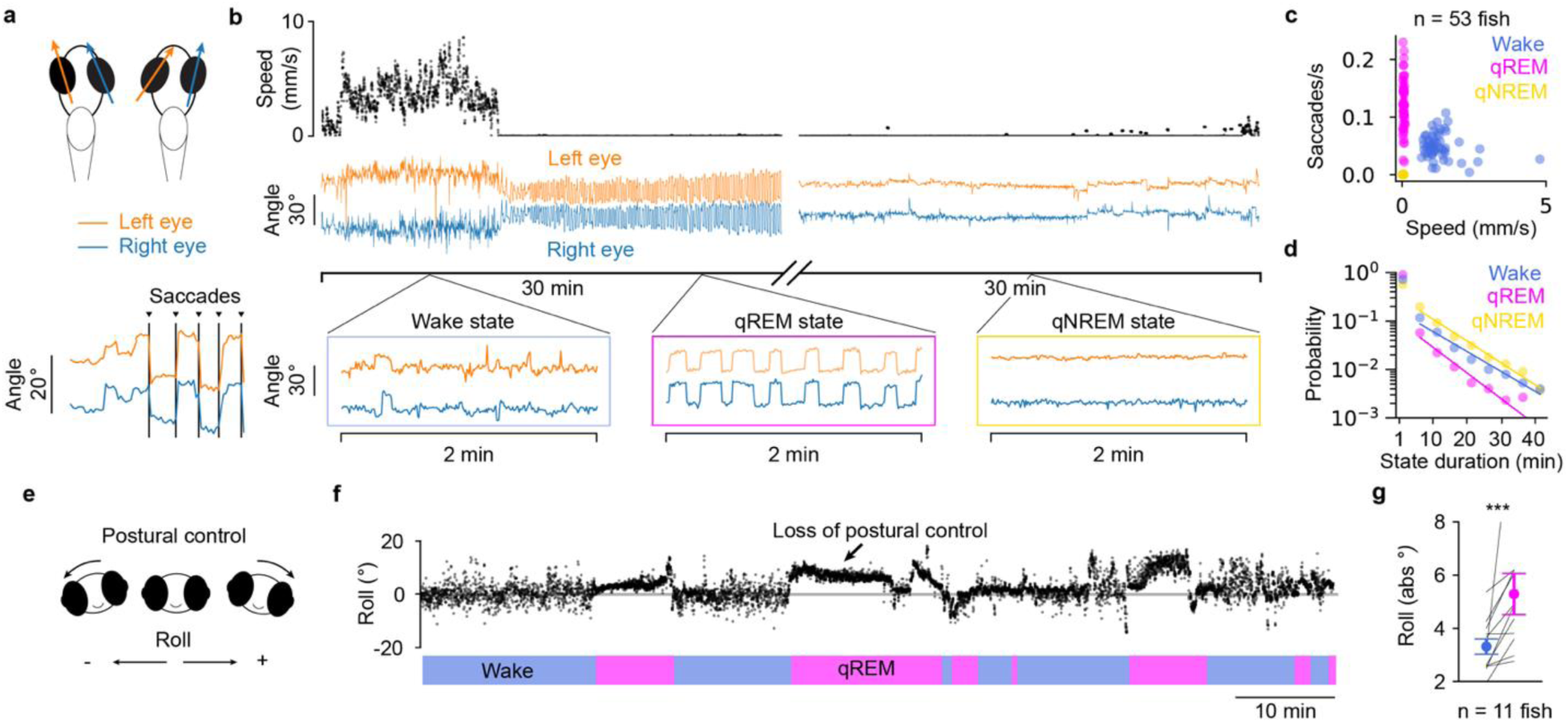
Identification of a novel, quiescent, sleep-like state in larval zebrafish. **a,** Measurement of fish eye angle. Eye angle is defined as the angle between the major axis of each eye and the animal heading. Saccades were identified as large amplitude, simultaneous deflection of both eyes in the same direction (**Methods**). **b,** Two 30 min example excerpts from a representative experiment. Insets of eye angles illustrate that quiescence can be subdivided based on the distinct eye movement signatures in the qREM and qNREM states (quiescence-REM vs. quiescence-NREM). **c,** A hierarchical HMM was used to classify each time point from all experiments as one of three distinct behavioral states. Inputs to the model were fish speed and eye saccade rate. Scatter plot shows the mean of speed and saccade rate per animal for each HMM-labeled behavioral state (n = 53 fish, 24 hr recording with 14 hr light / 10 hr dark, **Methods**). **d,** Duration probability histogram for each behavioral state plotted in 5 min bins. Lines represent least squares exponential model fits to underlying data. **e,** Roll was defined as the degree of head rotation about the animal’s rostral-caudal axis (0 degrees corresponds to normal upright posture, **Methods**). **f,** Example experiment showing the distribution of roll for a single animal over a 90 min period. **g,** The mean absolute roll was calculated for each behavior state. Large values indicate a large average deviation from normal upright posture. Roll was significantly higher during qREM compared to wake (*p =* 0.001, n = 11 fish, Wilcoxon signed rank

Strikingly, quiescence could be subdivided into two distinct states based on eye movements. The first state contained minimal eye movement (qNREM), consistent with observations from immobilized animals after sleep deprivation^28^, while the second contained high amplitude, regular eye saccades (**Fig. 1b**). We fit a 3-state hierarchical HMM to longitudinal measurements of speed and saccade rate (**Extended Data Fig. 1, Methods: Hierarchical HMM**). The three resulting behavioral states, waking non-quiescent state (wake), quiescence with Rapid Eye Movements (qREM), and quiescence with No Rapid Eye Movements (qNREM), were clearly separated along the speed and saccade rate axes (**Fig. 1c, Supplementary Video 1**). In addition, all three states were stable and long-lived (state durations of 9.3 ± 31.0 min, 3.4 ± 5.8 min, 8.9 ± 12.0 min for wake, qREM, and qNREM states respectively, mean ± s.d., n = 53 fish, **Fig. 1d**). The HMM output was enriched for relatively short state durations, but the probability of state durations lasting longer than 5 minutes followed an exponential distribution (λ = wake: −0.10 min^-1^, qREM: −0.12 min^-1^, qNREM: −0.10 min^-1^, pooled distribution of n = 53 fish) (**Fig. 1d**).

Sleep in mammals, and in particular REM sleep, is associated with loss of postural control^38^. Loss of postural control has also been observed during sleep in adult fish^39^. Therefore, we next measured the roll and pitch of animals during both qREM and wake to determine if loss of postural control was associated with the qREM state (**Fig. 1e-g, Methods**). While pitch was not significantly different between wake and qREM (p > 0.05, n = 11 fish, Wilcoxon signed rank test), roll showed clear state-dependent changes (**Fig. 1f-g**). During wake, roll was distributed evenly around zero (**Fig. 1f**). However, during qREM, roll was significantly skewed (either positively or negatively) for the duration of the state (mean absolute roll wake: 3.31 ± 0.29 vs. qREM: 5.29 ± 0.77°, mean ± s.e., *p =* 0.010, n = 11 fish, Wilcoxon signed rank test, **Fig. 1g**). Thus, qREM is associated with a distinct loss of postural control, reminiscent of a sleep-like state.

### qREM is associated with an increased arousal threshold

In addition to reduced locomotion and loss of postural control, a hallmark of sleep is increased arousal threshold^3,4^. To determine if arousal threshold is modulated during the qREM state, we performed arousal assays in two distinct sensory modalities (**Fig. 2**). First, we presented fish with a series of “dark flash” visual stimuli. Whole field dark flashes are known to elicit large amplitude startle responses in larval fish^40^ (**Fig. 2a, Methods**) which are reduced during sleep^3^. We found high startle response probability during wake (0.58 ± 0.05, mean ± s.e.) and significantly suppressed response probability during qREM (0.14 ± 0.03, p = 4.2e-7, n = 24 fish, Wilcoxon signed rank test, **Fig. 2b-c**), indicating an increase in arousal threshold during qREM. By definition, movement levels were at or near zero during the qREM state. Therefore, we performed a control analysis to ensure that the difference in response probability between wake and qREM could not be statistically explained by changes in overall movement levels between the two states (**Extended Data Fig. 2a-c**). After controlling for changes in gross motor movement between the two states, response probability remained reduced, confirming that arousal threshold is increased during qREM.

**Figure 2.**
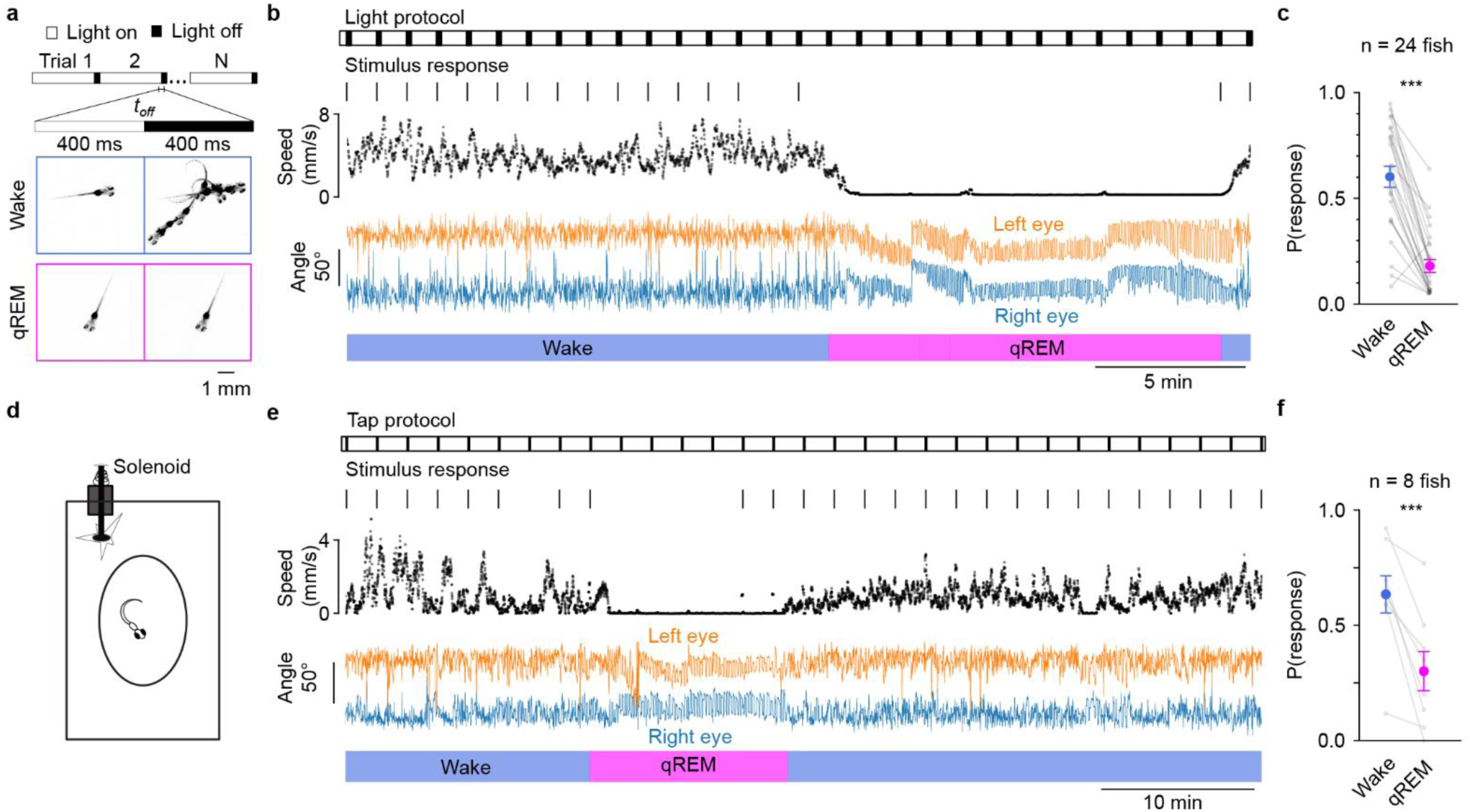
qREM is associated with an increased arousal threshold. **a,** Top: Experimental protocol for dark flash experiments. Each trial consisted of a 1 s whole field dark flash stimulus with an inter-trial interval of 60 s. Bottom: Representative example trial during wake (top) and during qREM (bottom). In each of the four panels, a subsample of 10 fish images spanning 400 ms are overlaid. Center line divides time points before (left) and after (right) light offset. **b,** 30 min example data excerpt from a dark flash experiment. Dark flash stimuli reliably elicited high amplitude startle responses (**Methods**) during wake, but not during qREM. **c,** Startle response probability as a function of behavior state. Responses are significantly more likely during the wake state (*p* = 4.2e-7, n = 24 fish, Wilcoxon signed rank test). **d,** Experimental setup for mechano-acoustic tapping experiments. Every 2 min, a solenoid struck the plate that the fish’s chamber was mounted on, eliciting a robust startle response. **e,** 60 min example data excerpt from a tapping experiment. Tap stimuli reliably elicited startle responses during wake (0.63 ± 0.08, mean ± s.e.), but not during qREM (0.30 ± 0.08, mean ± s.e.). **f,** Startle response probability as a function of behavior state. Responses were significantly more likely during wake (*p* = 0.008, n = 8 fish, Wilcoxon signed rank test).

Second, we tested whether sensory modalities other than vision were also affected by qREM. To do this, we performed a mechano-acoustic tapping assay, which, like dark flashes, elicits startle responses in alert fish^41^ (**Fig. 2d, Methods**), and has been used to quantify arousal levels across sleep and wake ^42,43^. Again, response probability to mechano-acoustic taps was significantly reduced during the qREM state (wake: 0.63 ± 0.08, qREM: 0.30 ± 0.08, mean ± s.e, *p* = 0.008, n = 8 fish, Wilcoxon signed rank test; **Fig. 2 e-f**) and this effect could not be explained by the decreased overall movement levels during quiescence (**Extended Data Fig. 2d-f**). Thus, qREM states are associated with the suppression of sensory systems across modalities. Collectively, these experiments establish that qREM has features of a sleep-like state, with prolonged immobility (**Fig. 1b-c**), loss of postural control (**Fig. 1e-g**), and increased arousal threshold (**Fig. 2**).

### Circadian and homeostatic regulation of qREM

We next asked how the expression of qREM is organized throughout the 24-hour circadian cycle. Using a high-throughput, high-resolution (50 µm/px, 50 frame/s), wide field of view (FOV, 207 x 179 mm) imaging system, we simultaneously monitored the activity of multiple animals (up to 20 fish per experiment) over the course of 24 hours (**Fig. 3a-c**, **Methods**). Lighting conditions were matched to the 14 / 10 hr light / dark schedule that fish were reared in. Surprisingly, qREM was almost exclusively restricted to circadian daytime hours and qNREM to nighttime (**Fig. 3d-e**). The probability of observing qREM during the day was 0.36 ± 0.03 while the probability of observing qREM during the night was only 0.05 ± 0.01 (mean ± s.e., *p* = 1.2e-9, n = 53 fish, Wilcoxon signed rank test, **Fig. 3e**). Conversely, the probability of observing qNREM was 0.02 ± 0.01 during the day but was 0.83 ± 0.01 during the night (mean ± s.e., *p* = 2.4e-10, n = 53 fish, Wilcoxon signed rank test, **Fig. 3e**, **Extended Data Fig. 3a**).

**Figure 3.**
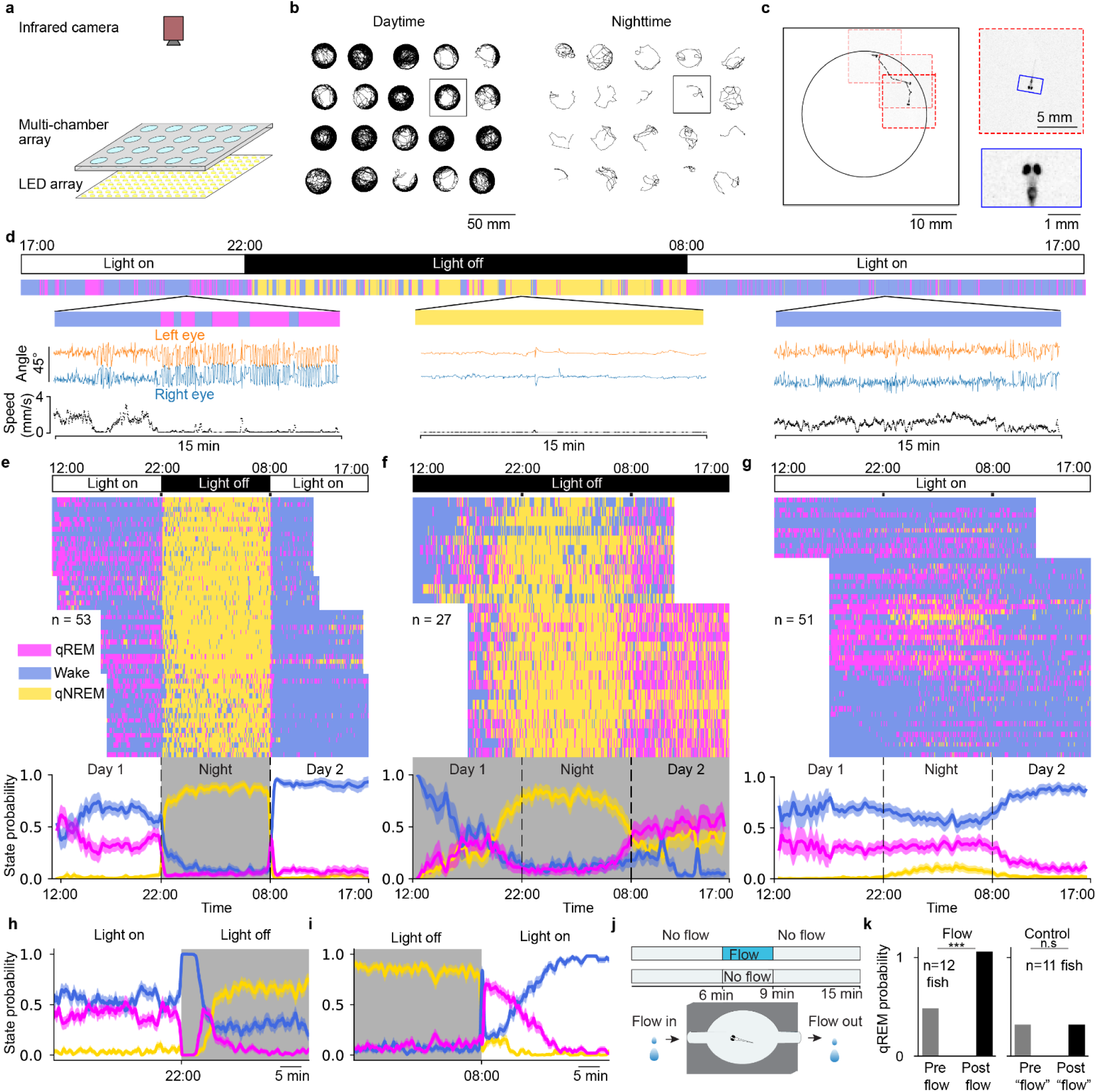
Circadian and homeostatic regulation of qREM. **a,** Experimental setup for longitudinal zebrafish behavioral recordings. **b,** Example zebrafish movement trajectories during 1 hour of daytime (left) and nighttime (right). Each fish was housed individually in a 28 mm × 28 mm × 1 mm chamber. **c,** Live tracking (50 frames/s) of zebrafish at high spatial resolution (50 µm / pixel) allowed detailed measurements of eye angle. Red dashed boxes represent the actively tracked region of the camera sensor (256 × 256 pixels) which moves with the fish location to enable high speed tracking and data reduction. **d,** One example dataset recorded using the experimental setup in **a**. On top are the light conditions which are matched to the circadian cycle of the zebrafish housing facility. Below are the HMM-inferred behavioral states, qREM (magenta), qNREM (gold) and wake (blue). Insets highlight eye angles and speed. **e-g,** Raster plot (top) and probability (bottom) of HMM-inferred behavioral states during light cycled (**e**, n = 53 fish, 14 hr light /10 hr dark), constant dark (**f,** n = 27 fish), and constant light (**g,** n = 51 fish) conditions. **h,** Probability of each behavioral state at the transition from light to dark (a zoom of light-dark transition from **e**). **i,** Same as **h** for dark to light transitions. **j,** Experimental setup and protocol for flow experiments. Following a pre-flow baseline period, E3 water is flowed into the chamber (**Methods**) for 3 minutes. In control experiments there was no flow. **k,** Probability of observing qREM in pre-flow (6 min) vs. post-flow (6 min) periods in experiments with flow (left) and in control experiments without flow (right).

To determine if the temporal segregation of qREM and qNREM was driven by environmental luminosity (i.e., light vs. dark) or by internal circadian rhythm (i.e., time of day), we performed two additional sets of experiments: 24 hours of constant dark (**Fig. 3f**) and 24 hours of constant light (**Fig. 3g**).

Experiments during constant luminosity conditions revealed that qREM and qNREM occupancy are regulated by an internal circadian clock. In constant dark, qREM probability was higher during the day and lower during the night (0.26 ± 0.02 daytime vs. 0.12 ± 0.01 nighttime, mean ± s.e., *p* = 1.0e-7, Wilcoxon signed rank test, n *=* 27 fish). Conversely, qNREM probability was higher during circadian night and lower during circadian day (0.36 ± 0.03 daytime vs. 0.76 ± 0.02 nighttime, mean ± s.e., *p* = 1.5e-8, Wilcoxon signed rank test, n = 27 fish, **Fig. 3f**), and followed a similar pattern in constant light conditions (qNREM probability: 0.01 ± 0.01 daytime vs. 0.08 ± 0.02 nighttime, mean ± s.e., *p* = 1.3e-9, Wilcoxon signed rank test, n = 51 fish, **Fig. 3g**).

qREM and qNREM probabilities are biased by environmental luminosity as well. During circadian night, qREM was more probable in the light than in the dark (light on: 0.32 ± 0.04, n = 51 fish vs. light off: 0.05 ± 0.01, n = 53 fish, mean ± s.e., *p* = 1.4e-9, rank sum test) while, conversely, qNREM was less likely in the light (light on: 0.08 ± 0.02, n = 51 fish vs. light off: 0.83 ± 0.01, n = 53 fish, mean ± s.e., *p* = 1.6e-18, rank sum test). Collectively, these findings argue that qREM and qNREM are governed by a combination of internal circadian timing and external environmental luminosity.

We found that environmental luminosity could also modulate qREM occupancy on faster timescales. Both the transitions from light to dark, and from dark to light, triggered a distinct sequence of state transitions (**Fig. 3h-i**). In agreement with prior work^40^, light to dark transitions led to a stable period of high locomotor activity which lasted several minutes. Interestingly, this was followed immediately by a qREM rebound (**Fig. 3h**). At the transition from dark to light, we also observed a transient spike in high locomotor activity immediately followed by a rebound in qREM for many minutes (**Fig. 3i**).

Finally, we asked whether the previous experience of the animal and accumulated homeostatic pressure can lead to a subsequent rebound in qREM^44^. Recent work in zebrafish has demonstrated that accumulation of homeostatic pressure due to increased global brain-wide activity leads to an immediate rebound in sleep occupancy^45^. To determine whether this rebound primarily consists of qREM or qNREM sleep, we manipulated global brain activity using water flow as a naturalistic stimulus (**Fig. 3j-k**, **Extended Data Fig. 4**), which has previously been shown to trigger sleep rebound^46^. We found that water flow is associated with a sudden increase in global brain activity (**Extended Data Fig. 4**). During 1 min of constant flow, 62 % of neurons increased their activity relative to pre-flow baseline activity (n = 10 fish, **Extended Data Fig. 4b, Methods: Water flow experiments).** After termination of the flow stimulus, the probability of qREM increased significantly during the six-minute window post flow (pre-flow qREM probability: 0.42 vs post-flow probability: 0.92, *p* = 0.0004, n = 12 fish, Binomial Test, **Fig. 3k**). Thus, naturally induced increases in global brain activity are correlated with subsequent rebound in qREM.

**Figure 4.**
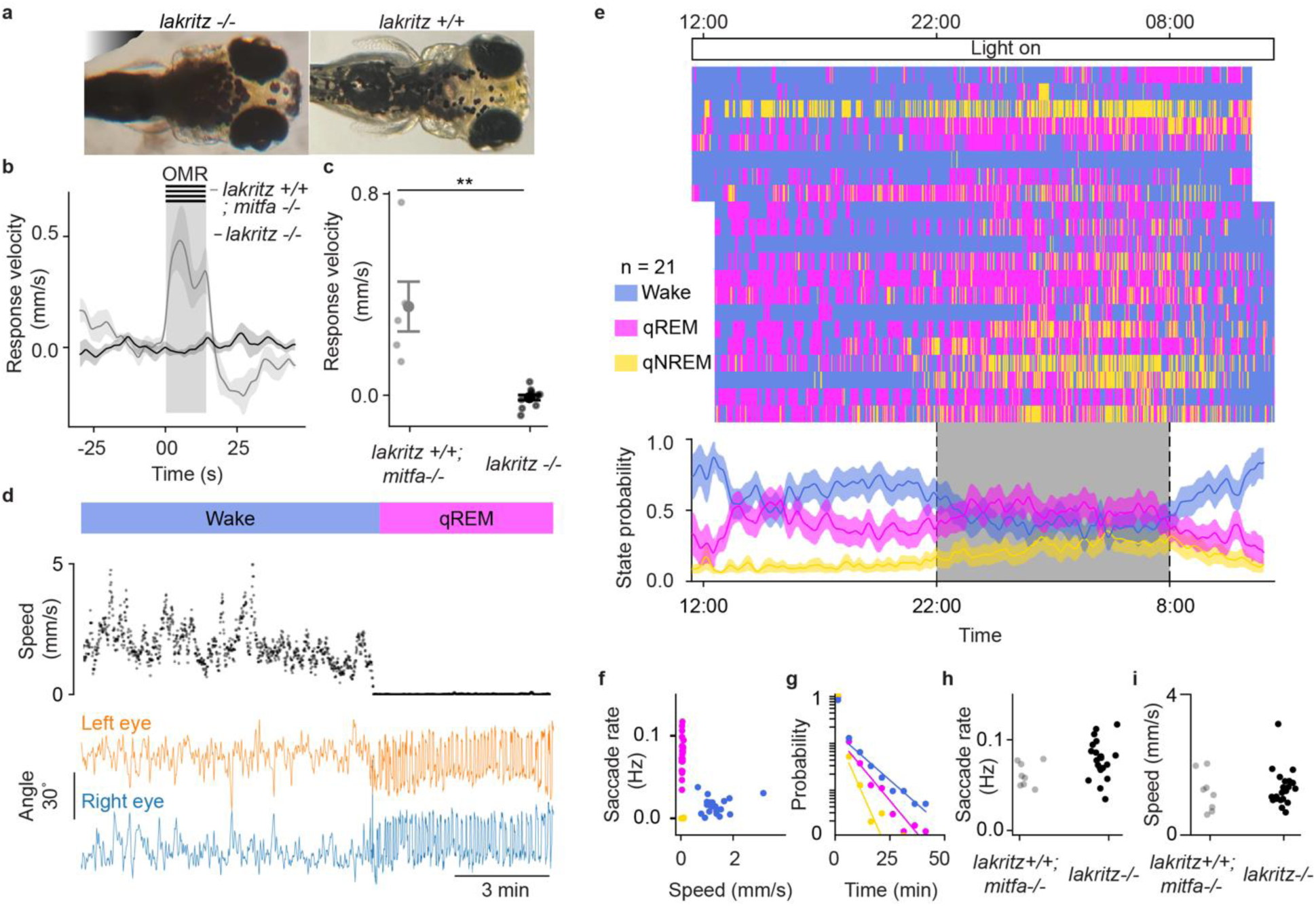
qREM in congenitally blind (*lakritz −/−*) zebrafish. **a**, Hyperpigmented *lakritz* −/− animal (left) and wildtype *lakritz +/+* animal (right). **b**, Optomotor response (OMR) of *lakritz* −/− and *lakritz+/+; mitfa−/−* animals to moving grating. Black and grey traces are smoothed average response velocities (σ = 1 s) in the direction of moving gratings for *lakritz −/−* (black, n = 11 fish) and *lakritz +/+*; *mitfa −/−* (grey, n = 5 fish) animals, respectively. The shaded region from 0-15 s represents the OMR stimulus period. The inter-stimulus interval was 45 s. **c**, OMR response velocity summary for *lakritz* −/− and *lakritz +/+; mitfa −/−* animals. **d**, Example speed (middle) and eye angles (bottom) of a *lakritz* −/− animal over 15 min, with inferred HMM state labels (top). **e**, Raster plot (top) and probability (bottom) of HMM-inferred behavioral states of *lakritz* −/− animals in constant light conditions. The shaded period in the bottom represents circadian night. **f**, Scatter plot of mean speed and saccade rate of *lakritz* −/− animals for each HMM-inferred behavioral state in constant light conditions. **g,** Duration histograms of *lakritz* −/− animals binned at 5 min for each behavioral state. **h**, Mean saccade rate per animal comparison between *lakritz* −/− animals (n = 21 fish) and condition-matched *lakritz +/+; mitfa −/−* controls. **i,** Comparison of mean speed between *lakritz* −/− and *lakritz +/+; mitfa −/−* animals. n = 21 *lakritz* −/− animals and n = 8 *lakritz +/+; mitfa −/−* control animals (imaged in the same experimental session) were used for **e-i**.

### Eye movements during qREM are not dependent on visual feedback

The presence of qREM during constant darkness suggests that visual feedback is not necessary to maintain qREM. However, it remains possible that past visual experience is necessary for qREM to occur. To test this, we performed 24-hour experiments with *lakritz−/−* mutant fish which lack retinal ganglion cells^47^ and are congenitally blind. We identified *lakritz−/−* mutants by their characteristic hyperpigmentation (**Fig. 4a**) and confirmed the blindness of the hyperpigmented animals by their lack of responsiveness to visual stimuli (**Fig. 4b-c**). The optomotor response (OMR) velocity of the hyperpigmented animals was not significantly different than zero (−0.3 ± 0.02 mm/s, mean ± s.e, n = 11 fish) and was significantly lower than that of *lakritz+/+; mitfa−/−* controls (0.88 ± 0.24 mm/s, mean ± s.e., *p* = 1.8e-3, n = 5 fish, Wilcoxon rank sum test).

Remarkably, *lakritz−/−* mutants clearly expressed qREM states (**Fig. 4d**), suggesting that eye movements during qREM do not depend on prior visual experiences or current visual feedback. Across 24 hours, *lakritz−/−* mutants cycled through all three behavioral states (wake, qNREM, and qREM, **Fig. 4e**) which separated well along the saccade rate and speed axes (**Fig. 4f**). Compared to *lakritz+/+; mitfa−/−* controls, *lakritz−/−* mutants do not show a meaningful difference in their state durations (**Fig. 4g**), saccade rate (**Fig. 4h**, *lakritz−/−*: 0.077 ± 0.004 saccade/s, *lakritz+/+; mitfa−/−*: 0.061 ± 0.004 saccade/s, mean ± s.e., p = 0.045, Wilcoxon rank sum test), or overall movement levels (**Fig. 4i**, *lakritz −/−*: 1.32 ± 0.11 mm/s, *lakritz+/+; mitfa−/−*: 1.22 ± 0.18 mm/s, mean ± s.e., *p* = 0.732, Wilcoxon rank sum test).

Finally, in *lakritz−/−* mutants, wake state probability remained significantly higher during circadian day in comparison to circadian night (0.60 ± 0.04 daytime vs. 0.36 ± 0.05 nighttime, mean ± s.e., *p* = 9.5e-7, *n* = 21 fish, Wilcoxon signed rank test) and qNREM state probability was significantly higher during night than day (0.074 ± 0.020 daytime vs. 0.193 ± 0.039 nighttime, mean ± s.e., *p* = 9.5e-7, *n* = 21 fish). This suggests that internal circadian timing persists in the blind *lakritz−/−* mutant fish, possibly through intrinsically photosensitive structures such as the pineal gland^6,43^.

### Neural dynamics during qREM

To investigate the neural dynamics of qREM, we performed brain-wide calcium imaging in freely swimming animals expressing pan-neuronal H2B-GCaMP6s (5-8 dpf) while continuously recording both locomotion and eye movements^48^. Putative single neuron activity traces were extracted using Non-negative Matrix Factorization^49^ (NMF, **Methods**). Using the 3-state HMM described previously (**Extended Data Fig. 1**), we classified each time-point as belonging to wake, qREM, or qNREM state. By combining neural and behavioral imaging, we were able to identify three key neural features of qREM. First, qREM led to broad suppression of neural activity; second, distinct populations of neurons encoded the behavioral features of qREM (e.g., eye saccades) vs. the qREM state itself; and third, global brain dynamics evolved in an organized way in time across the qREM state.

To determine how overall brain activity changed between qREM and wake, we measured mean calcium activity during each state. During qREM, neural activity was broadly suppressed across the brain (**Fig. 5a, Supplementary Video 2**). Across fish (n = 11 fish), 73.7 ± 4.2 % (mean ± s.d.) of all neurons were negatively correlated with the qREM state. Using post-hoc *in situ* labeling of noradrenaline-expressing cells^50^, we found that the noradrenergic locus coeruleus was strongly suppressed during qREM (**Fig. 5b-d**), supporting our behavioral evidence that qREM is a low arousal sleep-like state^51,52^.

**Figure 5.**
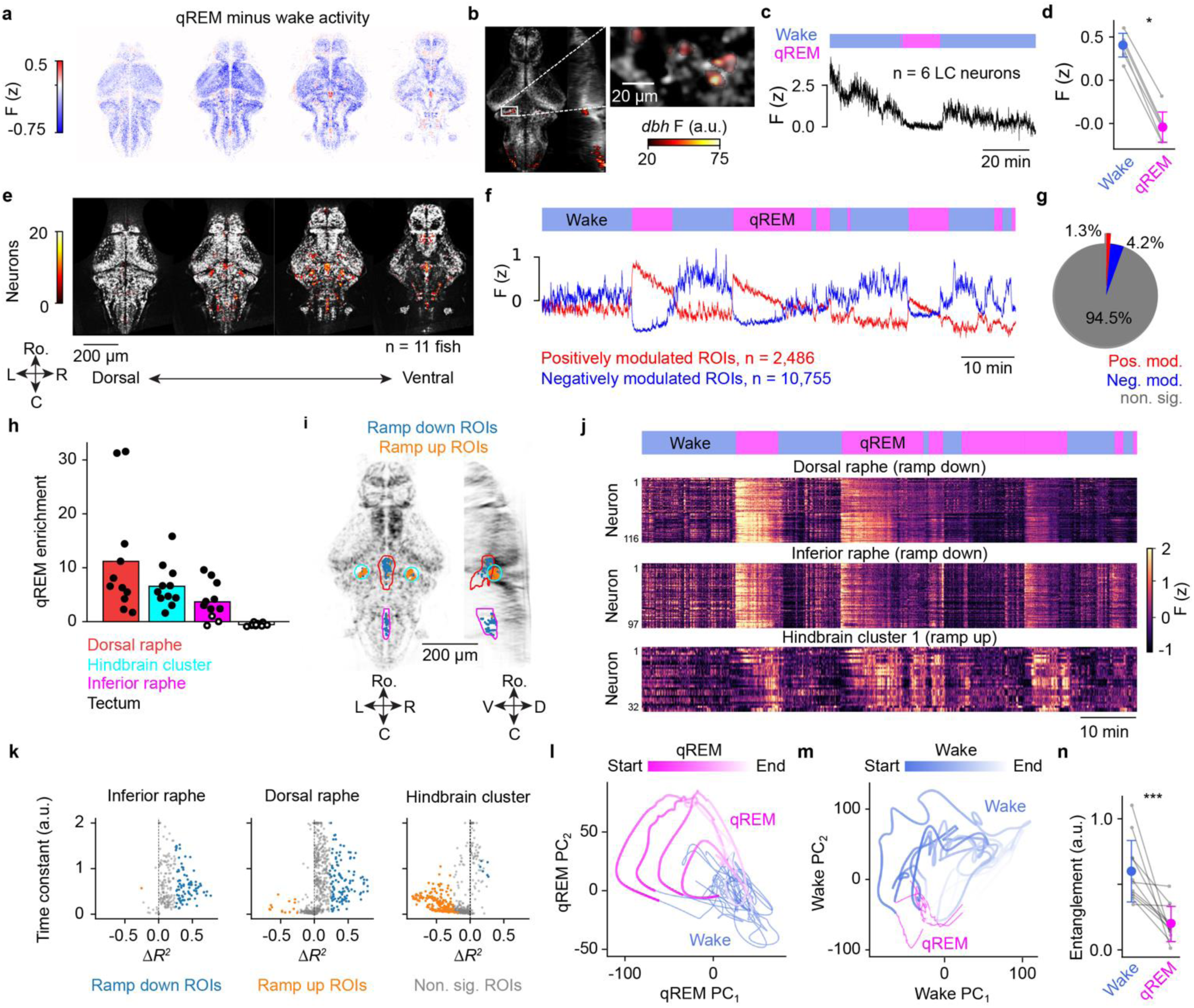
Whole-brain analysis reveals dynamic qREM state encoding brain networks. **a,** n = 11 fish were transformed into the mapZebrain atlas space (**Methods**). The coordinate space was binned and the mean qREM minus wake calcium activity across all neurons belonging to each bin is shown here. Data was split into 5 even quantiles, spanning the dorsal ventral axis of the reference brain, and the four most dorsal quantiles are shown here. **b,** Example reference brain image for one fish with overlaid colormap indicating the intensity of *in-situ* labeling of *dbh.* Inset highlights the left locus coeruleus (LC) of this animal. **c,** Mean of raw fluorescence activity measured across the six *dbh* positive LC cells shown in panel **b**. **d,** Mean calcium activity during wake (blue) and qREM (magenta) across all fish in which post hoc *dbh* labeling was performed. Each grey line represents mean across neurons in one fish. Activity was significantly higher during wake than during qREM (*p* = 0.03, n = 6 fish, Wilcoxon signed rank test). **e,** The dorso-ventral axis was divided as in panel **a**. Heatmap intensity corresponds to the number of positively modulated qREM neurons at each location across all fish (n = 11 fish). **f,** Mean activity across positively modulated (red) and negatively modulated (blue) neurons in one example fish. **g,** Fraction of neurons that were not significantly modulated during qREM (grey), negatively modulated qREM (blue), or positively modulated qREM (red) across all fish. **h,** Fold-enrichment of positively modulated neurons in each brain region relative to a random anatomical sampling of neurons (**Methods**). Each dot represents the enrichment per region for one fish (filled dots: *p* < 0.05) and bars represent the mean across fish. **i,** Anatomical organization of positively modulated qREM neurons belonging to the 3 brain regions with significant enrichment highlighted in panel **h. j,** Activity rasters for one example fish in the three brain regions highlighted in **i. k,** Exponential model fits for positively modulated neurons (**Methods**). Each dot represents one neuron. Neurons from all fish are included (*n* = 11 fish). Neurons were categorized into three groups, ramp up (orange), ramp down (blue), and non-ramping (grey). Δ*R^2^* refers to *R^2^* of exponential decay model minus *R^2^* of “1 minus exponential” ramp up model fits. **l,** Whole-brain PCA performed only during qREM periods. All timepoints are projected onto the top two Principal Components (PCs) of this space and colored by qREM (magenta) vs. wake (blue). Colormap corresponds to time within each respective qREM period. **m**, Same as **l,** with PCA performed on wake periods only. **n**, Mean entanglement of wake (blue) vs. qREM (magenta) PCA trajectories for each fish. Wake trajectories were significantly more entangled (*p* = 0.001, n = 11 fish, Wilcoxon signed rank test). Entanglement was defined as the number of self-crossings within a single state trajectory, normalized by the length of the state (**Methods**).

Despite broad suppression of neural activity, a subset of brain regions were significantly more active during qREM than during wake (**Fig. 5e**). Neurons in these brain regions could encode either the characteristic motor patterns of qREM (e.g., eye saccades), or the qREM state itself. To distinguish between these two possibilities, the activity of each neuron was regressed against both behavioral and qREM state variables using multiple linear regression (**Methods: Multivariate Linear Regression Analysis**). To determine which aspects of behavior and / or internal state were encoded by single neurons, we compared the full model performance (*R^2^*) to a set of additional partial models in which specific subsets of variables were excluded (**Methods: Multivariate Linear Regression Analysis**). A regression variable contributed significantly to a neuron’s activity if the full model significantly outperformed the partial model.

We found that neurons encoding qREM-associated eye movements were primarily located in the abducens nucleus, consistent with prior work^53^ (**Extended Data Fig. 5a-b**). Unlike previous studies in immobilized animals^54^, we found a clear anatomical segregation of saccade encoding neurons from neurons encoding left and right turn bias (**Extended Data Fig. 5b**). In waking states, locomotor neurons were active, while saccade neurons were largely suppressed (**Extended Data Fig. 5c-d**). This is consistent with our behavioral observation that large amplitude, stable eye saccade events are mostly decoupled from swimming movements (**Fig. 1b-c**, **Extended Data Fig. 5c-d**).

In addition to neurons encoding behavioral features of qREM, we identified 991 ± 720 (mean ± s.d., n = 11 fish) neurons per fish that uniquely and positively encoded the qREM state itself (**Fig. 5e-g**). These putative qREM neurons were concentrated primarily in the ventral hindbrain and made up 1.3 % of all recorded neurons, on average (**Fig. 5g**). qREM-encoding cells were significantly enriched in the dorsal and inferior raphe, as well as in an adjacent lateral hindbrain cluster (fold enrichment in dorsal raphe: 12.2 ± 10.6, inferior raphe: 4.8 ± 3.5, lateral hindbrain cluster: 7.6 ± 4.0, mean ± s.d.) (**Fig. 5h, Methods: Anatomical enrichment**).

Strikingly, the activity of many qREM state-encoding neurons was not a simple binary indicator function of behavioral state. Instead, they exhibited both positive and negative ramping dynamics that spanned the duration of each qREM period (**Fig. 5i-k**). We fit exponential curves to each neuron’s qREM activity to quantify the dynamics of these ramps (**Methods: Exponential Model Fits to qREM Activity**). Ramping shapes were stereotyped according to anatomical location (**Fig. 5i-k**). In both the inferior and dorsal raphe, most neurons (i.e., the “ramp down neurons”) were best fit by exponential decay and decay time constants were significantly larger in the dorsal raphe than in the inferior raphe (dorsal raphe: 0.91 ± 0.03, inferior raphe: 0.67 ± 0.03, mean ± s.e., *p* = 1.3e-6, rank sum test; duration of each REM period was rescaled to range from 0 to 1). Conversely, lateral hindbrain cluster neurons were best fit by “1 minus exponential” dynamics (ramp up neurons) with relatively short time constants (0.21 ± 0.01, mean ± s.e.). Notably, the ramp up neurons were not as well fit by the exponential models in general (*R^2^* for hindbrain ramp up neurons: 0.05 ± 0.06 vs. dorsal raphe neurons: 0.17 ± 0.16 and inferior raphe neurons: 0.23 ± 0.17, mean ± s.d.), suggesting that their activity patterns are more complex. Indeed, for many neurons in this region, activity did not seem to smoothly ramp up so much as the probability of becoming active increased as a function of qREM time elapsed (**Fig. 5j**).

Given the nontrivial encoding of qREM state by individual neurons, we performed whole-brain PCA to gain a more comprehensive view of global dynamics during qREM (**Methods: PCA of whole brain activity**). Contrary to the fixed-point attractor model of quiescence reported in *C. elegans*^15^, we observed substantial variance within qREM itself. To investigate this, we restricted our whole-brain PCA analysis to qREM timepoints only (**Fig. 5l, Methods**). In the resulting qREM-PC space, activity evolved in an organized fashion throughout time during qREM (**Fig. 5l**). This was consistent across fish (**Extended Data Fig. 6a**). Performing the same analysis on wake periods, we found qualitatively similar results (**Fig. 5m**, **Extended Data Fig. 6b**). However, wake trajectories appeared to be less organized, perhaps reflecting the diversity of the waking brain state. Rather than exhibiting a distinctive geometry, wake state-space trajectories were often overlapping (i.e., “tangled”^55^). Temporal entanglement was significantly lower during qREM than during wake (**Fig. 5n**, *p* = 0.001, n = 11 fish, Wilcoxon signed rank test, **Methods: Entanglement of state space trajectories**). This suggests that the highest variance dimensions of qREM activity can be used to decode time progression, while the highest variance dimensions of wake activity also reflect other variables in addition to internal state, such as motor activity. Thus, we conclude that neural activity during the qREM state is primarily dominated by neuronal populations with slow ramping dynamics. Collectively, these populations produce disentangled representations of time in the population activity state space.

**Figure 6.**
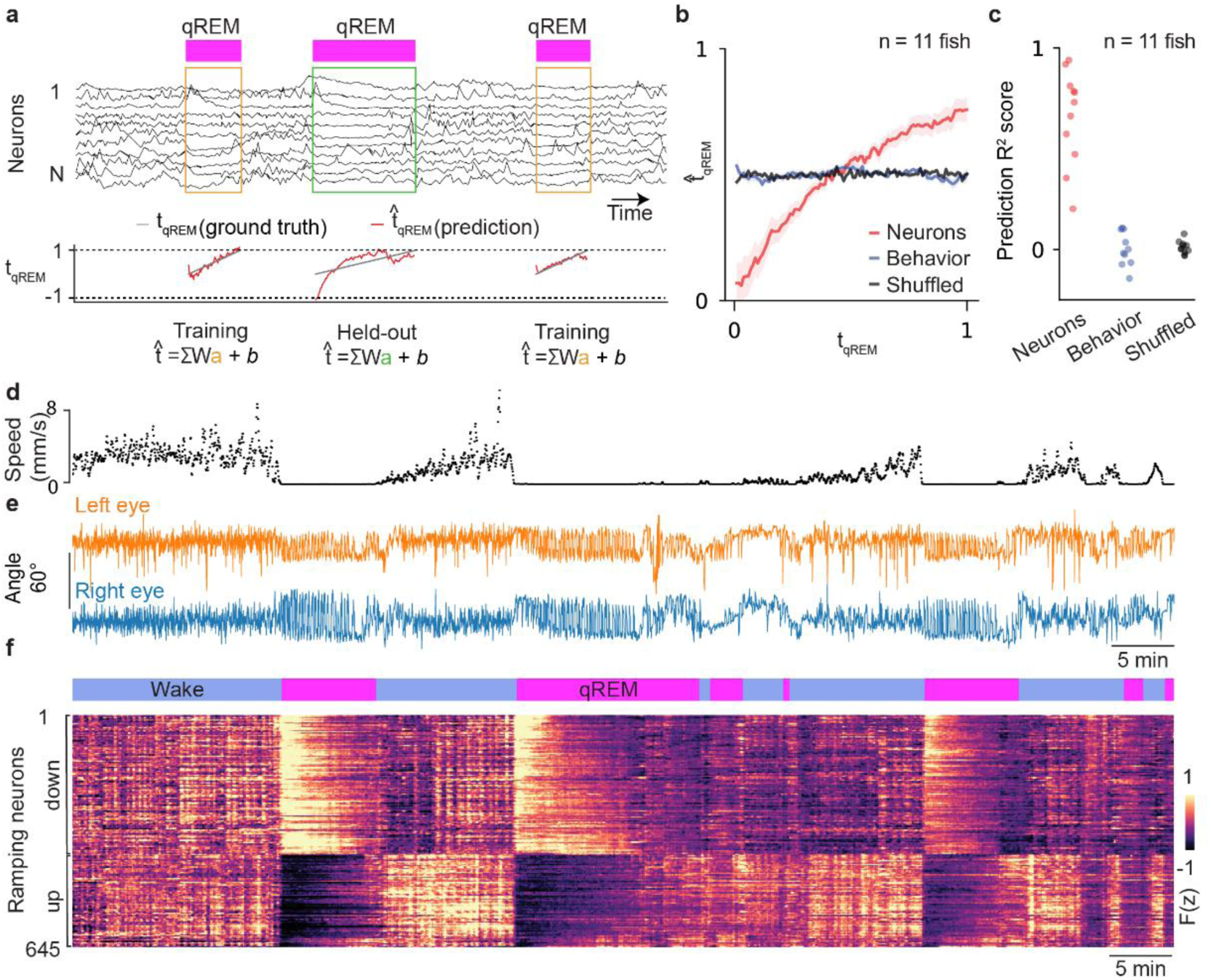
Decoding relative time progression in qREM from brain-wide neural dynamics. **a**, Relative time decoding procedure. Top: qREM state labels and activity of 11 example neurons smoothed with a Gaussian kernel (σ = 25 s). The linear decoder was trained on all qREM states except for one held-out period. Trained decoder weights were then applied to the held-out qREM period to predict relative time. This procedure was repeated for each qREM state in order to generate predictions for the entire dataset. Bottom: Relative time prediction (red) in each qREM state compared with true relative time (grey). **b**, Relative time prediction (mean ± s.e.) using only neural activity (red), behavioral variables (blue), or shuffled neural activity (black) (n = 11 fish). **c**. Decoder performance was quantified by measuring *R^2^* between true relative time and predicted relative time (neural *R^2^* = 0.66 ± 0.07, shuffled *R^2^* = 0.01 ± 0.01, behavior *R^2^* = −0.03 ± 0.04, mean ± s.e., n = 11 fish). **d-e,** Speed (mm/s) (**d**), left eye (**e**, orange), and right eye (**e**, blue) angle of an example freely swimming larva. **f**, Top: Behavioral state labels (blue: wake, magenta: qREM) throughout the experiment. Bottom: Raster map of z-scored neural activity for neurons in the example animal that significantly contributed to relative time decoding during qREM state: ramp-down neurons (388 neurons, upper rows), which had negative decoding weights, and ramp-up neurons (257 neurons, bottom rows), which had positive decoding weights.

### qREM state evolves in time according to stereotyped neural dynamics

The whole-brain PCA results (**Fig. 5l-m**) demonstrated that neural activity evolves stereotypically during qREM state. This insight motivated us to try decoding time progression in the qREM state from neural activity. To this end, we trained a regularized linear decoder (**Extended Data Fig. 7a-b, Methods: Relative time decoding from whole-brain activity**) to predict relative time within the qREM state using whole-brain neural activity (**Fig. 6a**). To avoid overfitting, we evaluated decoder predictions on held-out qREM states. Therefore, we restricted our analysis to datasets with at least two qREM periods (**Methods**). Neural activity predicted relative time progression through the qREM state accurately (**Fig. 6b-c**; neurons *R^2^*score: 0.66 ± 0.07, shuffled *R^2^*score: 0.01 ± 0.01, mean ± s.e., *p* = 9.7e-4, Wilcoxon signed rank test, n = 11 fish). The decoder utilized both ramp up and ramp down neural populations, which were enriched among qREM state encoding neurons (**Fig. 6d-f**, **Extended Data Fig. 7c-d, Methods**).

It is possible that the predictive neural activity might actually reflect subtle changes in behavior across time (e.g., saccade rate, roll, pitch, speed, eye angles), instead of internal state. To test this possibility, we trained an additional decoder to predict relative time using behavioral metrics alone (**Methods**). Unlike neural activity, behavior alone was not able to predict relative time in qREM (**Fig. 6b-c**, behavior *R^2^* score: −0.03 ± 0.04, mean ± s.e., n = 11 fish). This confirms that neural dynamics evolve in time according to the progression of internal state variables and are not simply a reflection of observable motor patterns.

Relative time decoding during qREM motivated us to investigate whether there are also latent dimensions of wakeful neural activity that encode time progression during wake state. To test this, we employed the same linear decoding strategy to decode relative time progression in the wake state (**Extended Data Fig. 8a**). Although decoding performance was slightly lower on average than for qREM, time progression during wake could also be predicted from neural activity alone (**Extended Data Fig. 8b-c**, neurons *R^2^*score: 0.60 +/− 0.05, shuffled *R^2^* score: −0.01 +/− 0.01, mean +/− s.e., *p* = 9.7e-4, Wilcoxon signed rank test, n = 11 fish). Across fish, wake ramp-up and ramp-down neural activity contributed to time decoding (**Extended Data Fig. 8d-f**). Unlike qREM, time progression in wake, for some fish, could be decoded above chance levels using behavior alone (**Extended Data Fig. 8b-e**, *R^2^* score: 0.12 +/− 0.06, mean +/− s.e.).

Collectively, these results suggest that sleep and wake are not fixed points in state-space that transition stochastically back and forth but, instead, follow stereotypical dynamical trajectories that enable decoding of time within each state.

## DISCUSSION

Using longitudinal, high resolution, behavioral imaging of naturalistic wake and sleep-like states in larval zebrafish, we uncover a novel sub-state of quiescence (qREM) in teleost fish. This upends longstanding assumptions about the nature and organization of sleep states in early vertebrates. For nearly two decades, sleep in larval zebrafish, as well as in *Drosophila* and *C. elegans,* has been defined primarily as periods of locomotor quiescence with increased arousal threshold^1–6^. This definition has led to convincing demonstrations of evolutionarily conserved sleep-regulating molecules across vertebrate species and provided vital insights into the neuromodulatory populations that promote and inhibit sleep^3,4,6,42,43,45,51,56,57^. In addition, a recent study in zebrafish found brain signatures similar to slow wave sleep (NREM / SWS) and paradoxical sleep (potentially analogous to REM) that could be observed upon extended sleep deprivation and pharmacological intervention^18^. However, these brain states were observed with the striking omission of eye movements, which are the key behavioral signature of REM sleep. Thus, two possibilities exist: REM sleep with eye movement emerged later in vertebrate evolution, or REM-like sleep in fish exists but has yet to be identified.

In support of the second possibility, we find that coordinated eye movements are prevalent during periods of prolonged quiescence (qREM) and are also coupled with an elevated arousal threshold and the loss of postural control, typical hallmarks of sleep. Notably, qREM occurs regularly in naturally swimming fish which were not sleep deprived or pharmacologically manipulated, raising the possibility that naturally occurring state transitions between wake, qREM, and qNREM may be suppressed in tethered preparations, and/or by extended sleep deprivation or pharmacological intervention.

The function and evolution of rapid eye movements during sleep has been intensely debated since the discovery of eye saccades during sleep in 1953^17^. A wide array of hypotheses have been put forward that include 1) the vigilance hypothesis, which proposes that REM provides periodic increase in arousal and threat detection during sleep^18^, 2) the oculomotor innervation hypothesis, which proposes that REM is “a mechanism for the establishment of the neuromuscular pathways involved in voluntary conjugate eye movements in both phylogenesis and ontogenesis”^14^, and 3) the “scanning” hypothesis, which proposes that eye movements are induced by visual imagery during dreaming^13^.

In mammals, the vigilance hypothesis and scanning hypothesis have difficulty in accounting for the fact that REM occurs in utero^58^, in premature infants before eye opening^59^, and in visually deprived newborn primates^32^.

Consistent with these observations, we find that larval zebrafish eye movements during quiescence are not dependent on visual experience or visual feedback, as qREM states also occur during experiments performed in constant darkness and are also present in congenitally blind larval zebrafish. Collectively, these observations suggest that the saccades during qREM are not sustained by visual feedback (e.g., to serve a vigilance function) and are not driven by reactivated visual imagery during sleep (e.g., the “scanning” hypothesis). Rather, the ancestral form of REM in vertebrates may be most consistent with the oculomotor hypothesis, which proposes that REM sleep serves to stimulate the oculomotor system during prolonged periods of rest.

We note that the oculomotor and motor learning hypotheses do not exclude a possible role for qREM in the processing of waking experience (e.g., the emotional memory hypothesis^21^), as we find that qREM probability increases following intense waking experience (e.g., in response to water flow). This result is also consistent with recent work in larval zebrafish that used pharmacological activation of brain activity to pinpoint integrators of homeostatic pressure during wake, and convincingly demonstrated that brain-wide activation leads to sleep rebound during the day^45^. Though the study did not specifically examine eye movements during the sleep rebound, our results (**Fig. 3**) suggest that it is likely due to an increase in qREM.

Perhaps our most surprising behavioral finding is that qREM and qNREM are segregated by circadian time, preferentially occurring during circadian day or night, respectively. Unlike previous studies that have suggested that zebrafish eye movements are simply a function of ambient luminosity^29^, we find that this circadian segregation of qREM and qNREM holds in constant darkness or constant light, as well as in congenitally blind zebrafish. This finding highlights a potentially important evolutionary distinction between teleosts and mammals, as REM sleep in mammals typically follows NREM during one consolidated circadian period in adults^13^. However, mammalian infants transition directly between wake and REM and the overall organization of REM and NREM is substantially different from adults^59^. Future studies are required to determine whether the circadian segregation of qREM and qNREM in fish is specific to the larval stage or is retained into adulthood.

Having identified the behavioral signature of qREM in larval zebrafish, we leveraged our ability to record neural activity during naturally occurring qREM periods to uncover the neural dynamics underlying this novel state. Consistent with qREM being a low arousal, sleep-like state, we find that noradrenergic activity in the locus coeruleus, which is known to positively modulate arousal^51,52^, is strongly suppressed upon transition into qREM. In contrast, the serotonergic superior and inferior raphe regions are both positively modulated upon entry into qREM. This is unlike in mammals, where serotonergic signaling is anticorrelated with sleep^60,61^, but consistent with prior zebrafish studies that implicate serotonin being involved in regulating quiescent states^42^.

Finally, using dimensionality reduction to capture brain-wide neural dynamics, we show that the transition from wake to qREM in zebrafish sets off disentangled, dynamic trajectories in state space, in striking contrast to fixed point attractor models developed in invertebrate systems^15^. Unlike fixed point attractor models, dynamic trajectories enable us to decode temporal progression throughout the qREM state. More in-depth future studies across species will be needed to determine whether this represents an evolutionary difference between invertebrates and vertebrates, or whether dynamic trajectories can also be shown to exist in invertebrate sleep.

This work updates the conceptual framework for sleep in early vertebrates. Zebrafish sleep has been an exceedingly productive area of study for the last two decades. The existence of two clearly distinct substates, qREM and qNREM, offers an exciting new perspective with which to reexamine and reframe previous observations, assays, and manipulations in greater depth. Overall, our findings prompt us to reconsider longstanding assumptions about vertebrate sleep. Moreover, they shed new light on the evolutionary origin and function of REM sleep across the animal kingdom.

## METHODS

### Behavioral imaging at high spatiotemporal resolution across many animals

We developed a new behavioral acquisition system called DASHER (Dynamic Addressable System for High-speed and Enhanced Resolution), which combines real-time animal tracking with deep image sensor integration to achieve enhanced spatiotemporal resolution with a variety of commercial image sensors to significantly boost spatial and temporal resolution around the animal of interest (Robson et al, in preparation). Briefly, two versions were used for behavioral experiments, a version optimized for detailed behavioral imaging of a single animal (version 1) with a commodity USB3 camera, and a version optimized for detailed behavioral imaging of up to 20 animals (version 2) with a higher bandwidth 25GigE camera.

In version 1, we use a USB3 Blackfly S BFS-U3-51S5M-C (FLIR) with a Sony IMX250 sensor. We configure the image sensor to read out only a small region of interest (ROI) encompassing the animal and its immediate surroundings (e.g., static environment such as walls or dynamic elements such as prey), thus significantly increasing the imaging framerate without sacrificing the overall field of view (FOV) or the spatial resolution of the ROI around the animal. As the animal moves, DASHER performs a live update of the left and top readout offsets of the image sensor in the camera, such that the ROI is dynamically updated to remain centered on the animal. Animal tracking is adapted from previous publication^48^. By using DASHER to address only a small portion of the sensor, we increase framerate of the Blackfly S camera by several fold, allowing us to achieve an imaging speed of 250 Hz with a spatial resolution of 20 μm and a FOV of 10.24 mm × 10.24 mm (512 × 512 pixels). We use this version for behavioral recording in Fig. 2, 5 and 6 and Extended Data Fig. 2, 4, 5, 6, 7 and 8. Strobed near-infrared (NIR) and white light illumination was previously described^49^.

In version 2, we used a large sensor camera with high bandwidth 25GigE camera interface (Emergent Vision Technology HB-25000-SB, 5328 x 4608 at 50 frames per second) to simultaneously image up to 20 animals per experiment across 24 hr. Each animal is placed into a separate circular chamber (28 mm × 28 mm × 1 mm). An ROI (256 × 256 pixels, corresponding to 12.8 mm × 12.8 mm at 50 μm/pixel) around each animal was streamed to disk. A large rectangular PCB (280 × 240 mm) with interleaved NIR (SFH4650-Z, OSRAM) and white (GW PSLM31.FM OSRAM) LEDs was placed below the behavioral chamber. We use this version for behavioral recording in Fig. 1, 3, 4, and Extended Data Fig. 1 and 3.

### Eye-tracking

Each NIR zebrafish image was transformed into an ego-centric view from a geo-centric view using the head direction of the fish. In the ego-centric view of the fish, only a small region of interest (~0.9 mm × 0.9 mm) around the fish was retained for further processing. Next, we processed this region of interest with a difference-of-Gaussian filter (σ_1_ = ~0.1 mm, σ_2_ = 0.06 mm) to extract the eyes only and remove background noise in the image. We defined the *y* position of the eyes as the peak of the maximum projection of pixel intensities along the body axis. The *x* position of each eye was computed by projecting pixel intensities along the axis orthogonal to the body axis and fitting a bimodal Gaussian mixture model. After getting the center coordinates for the left and right eyes, we defined a rectangular mask around each eye and computed the central moments of each. From the central moments, we measured each eye angle.

### Hierarchical HMM

To determine the behavioral state of a zebrafish over time, we used a 2-stage hierarchical Hidden Markov Model (HMM, Extended Data Fig. 1). In the first step, we fit a 2-state HMM to the average speed, computed using a sliding window of 5 s, of multiple fish (n = 53 fish), dividing our data into periods of locomotor waking state and quiescence state. In the second step, we extracted only the quiescence periods and fit a second 2-state HMM to saccade rate, thus splitting quiescence periods into two sub-states, qREM and qNREM, based on the presence or absence of regular, conjugate eye movements (“eye saccades”). We scored a saccade event as simultaneous deflection of both eyes (> 22°) in either direction. After scoring saccade events, we computed saccade rate as the average number of saccade events per second in a running window of 90 s. HMM models were fit jointly to all animals by considering the behavior of each fish as an independent observation sequence. Model parameters were estimated by finding a local maximum of the joint likelihood function of multiple observations using an Expectation-Maximization (EM) type method^64^. Model parameters were estimated using the data from 24-hour light cycle experiments.

Maximizing model likelihood tended to result in an enrichment of transient, artificially short-lived states, presumably due to overfitting. In order to exclude these transient transitions, we smoothed the posterior probabilities of hidden states for each fish using a Gaussian filter with a kernel size of 45 s, and then used the Viterbi algorithm on the smoothed posterior probabilities to find the most likely hidden state sequence.

### Dark flash arousal assay

We delivered full-field dark flash stimuli that are known to elicit high amplitude escape turns. Each experiment lasted two hours. Full-field dark flash stimuli lasting 1 s each were presented by turning off all visible white light in the arena once every 60 s for a total of 120 stimuli per experiment. Tail tracking was performed (see below) and the angle of the tip of the tail, relative to animal heading, in the 1 s response window (time during light off) was analyzed. The first response window time bin with a relative tail angle of greater than 180 degrees was counted as a light response. Response probability was computed separately for each behavioral state, where behavioral state was defined using the hierarchical HMM described above.

### Tail tracking

Tail tracking was achieved using an optimization-based algorithm described previously^49^. Briefly, the algorithm created a high-resolution spline to trace out the curvature of the tail for each NIR image of the animal. The spline, consisting of 37 segments, each 100-μm long (5 pixels), was parameterized by local angle change. A nested grid search was used to fit the optimized tail spline onto the fish tail. The algorithm was designed to achieve robust tracking even in the presence of bright artifacts near the tail as previously described^49^. 28 of 37 tail segments were used for the dark flash response analysis described above as the remaining segments extended way beyond the tail length of the fish.

### Mechano-acoustic tap arousal assay

A solenoid was mounted to the plate holding the fish chamber. Every 2 minutes, the solenoid tapped the plate, providing mechano-acoustic stimulation to the fish through the coupling of the glass chamber to the metal plate. Experiments lasted between 2 hours and 5 hours. Startle responses to the acoustic tap were defined as moments where the fish’s instantaneous speed in the one second window following the tap stimulus exceeded 150 mm/s. Response probability was computed separately for each behavioral state, as determined by the hierarchical HMM described above.

### Water flow experiments

To test whether water flow periods can induce qREM, we recorded behavior using DASHER (version 1) while animals experienced water flow in a perfusion chamber. We used an elliptical PDMS chamber (30 mm × 20 mm). An initial 6 min behavioral baseline was recorded without water flow. Subsequently, water flow (~2.5 ml/min) was introduced for 3 minutes. Following the water flow period, behavior was recorded for an additional 6 min. Control experiments of equivalent duration were conducted without water flow. The qREM state probability during pre- and post-flow periods was compared to assess the effects of water flow on qREM induction.

Flow experiments were also performed on the tracking microscope in order to simultaneously record behavior and neural activity. In these experiments, an initial 6 min of baseline activity was recorded without water flow, followed by 1 min of water flow and then an additional 10 min without flow. For the analysis shown in Extended Data Fig. 4, neurons that were effectively sampled at < 1 Hz or that were inactive (fluorescence distribution with mean < median) during a 3 min window around flow (1 min of flow ± 1 min) were excluded.

### Optomotor response to moving gratings

To verify that *lakritz −/−* mutants were indeed blind, we tested their optomotor response to moving grating stimuli using DASHER (version 2). Fish were allowed to swim freely in 40 mm x 20 mm behavioral chambers. A projector (Optoma HD29H) was used to display a moving, full contrast (black/white) grating (spatial period 3-10 mm) for 15 s every 45 s over the course of a 1 hour experiment. Behavioral responses were quantified as Gaussian kernel smoothed (σ = 1 s) animal velocity in the direction of the moving grating during the 15 s stimulus presentation. In one hour, 100 % of control *lakritz +/+; mitfa−/−* fish (n = 5 fish) and 0 *lakrtiz −/−* mutants (n = 11 fish) responded to the grating above chance levels (Fig. 4c).

### Simultaneous behavioral and neural imaging of larval zebrafish

Fish were allowed to swim freely in a circular behavior arena (28 mm in diameter and ~1 mm in height) as described in previous work^49^. The chamber was illuminated with white light delivered either to all sides of the behavioral arena or from above. For behavioral tracking, the arena was illuminated by four custom NIR light strips with narrow angle, 850 nm LEDs (SFH4655-Z, Osram) that directed infrared light into the chamber.

To image neural activity, we conducted tracking DIFF microscopy experiments as previously described^62^ but with an updated motion cancellation system that utilizes a custom 3 axis motorized stage (Mohan et al, in preparation). We imaged brain-wide calcium activity within a volume of 1014 x 764 x 150 μm at cellular resolution in freely swimming fish expressing pan-neuronal H2B-GCaMP6s or H2B-GCaMP8s at 2 volumes/s. The details of this system have been described previously^48^.

### High-resolution offline registration of fluorescent brain volumes for each animal

We matched each fluorescence image from the tracking microscope for each animal to a high-resolution reference brain volume collected from the same animal. The registration pipeline, which includes a 3D rigid transformation and a non-rigid transformation, was described previously^48^. Briefly, each individual DIFF-sectioned brain slice (the moving image) is aligned to a plane within the reference brain volume with a 3D rigid transformation. Registration is then refined by locally adjusting a subdivided deformable surface within the reference volume using a regularized piecewise affine transform.

Parameters extracted from the affine registration step were used to compute fish pitch and roll during each time point (Fig. 1e-g).

### Non-negative matrix factorization and post processing

After image registration, we extracted raw fluorescence *F*(*t*) across time (*t*) for each neuron in the brain using non-negative matrix factorization^65^ (NMF, n = 11 fish, 114,435 ± 21,267 neuronal centroids, mean ± s.d., per fish). NMF decomposes the recorded spatiotemporal image data into two factors – one containing the spatial area of each neuron and another containing the time-varying raw fluorescence of each neuron. We applied spatially constrained NMF to our whole-brain datasets. NMF was performed independently for each axial section of a given brain volume. To avoid redundant neuronal centroids, we merged ROIs which were likely to represent repeated sampling of the same neuron across multiple axial planes. We did this by identifying and excluding ROIs which were 4 µm or less apart from each other in all the three dimensions and were also highly correlated (> 0.8 correlation coefficient).

Due to free swimming fish behavior, each neuron could remain unobserved in any given brain sweep (e.g., because it fell outside the axial scan range of a given sweep). Therefore, we only included neurons for further analysis whose raw fluorescence traces had at least 50 % available data, indicating that the neuron was tracked well over the course of the experiment (i.e., an overall sampling rate of at least once per second). After this step of preprocessing, we were left with 74,215 ± 23,457 (mean ± s.d.) neurons per fish. Missing values in the raw fluorescence traces were linearly interpolated for all analyses that relied on continuously available data traces (whole brain PCA, time decoding from neural activity).

Finally, to account for changes in fluorescence baseline due to photobleaching, we fit an exponential decay to the fluorescence timeseries of each neuron to estimate slow timescale bleaching. To correct for this, we subtracted the estimated bleaching from the neuron’s raw fluorescence.

### Registration to a common reference brain across animals

We registered our live imaging data (moving image) to a reference atlas zebrafish brain (mapZebrain atlas^63^) using CMTK^66^. Registration was performed in three steps. First, an initial affine transformation was performed by matching the center of mass of our live data with the atlas brain. Next, we fit a full affine transformation (translation, rotation, scale, and shear). Finally, a non-rigid warping was applied to the affine transformed brain which maximized normalized mutual information between our brain and the atlas brain. We optimized the parameters of this fit to preserve cell morphology while achieving the best possible global alignment between our brain and the atlas brain. After testing many different parameter combinations, we arrived at the following procedure for registration:

Step 1: cmtk make_initial_affine --centers-of-mass ATLAS.nii FLOATING.nii initial.xform

Step 2: cmtk registration --initial initial.xform --dofs 6,12 --ncc --exploration 50 --accuracy 3 -o affine.xform ATLAS.nii FLOATING.nii

Step 3: cmtk warp -o nonrigid.xform --nmi --jacobian-weight 0 --accurate -e 18 --grid-spacing 256 --energy-weight 1e-1 --refine 3 --coarsest 4 --ic-weight 0 --output-intermediate --accuracy 3 --initial affine.xform ATLAS.nii FLOATING.nii

The resulting nonrigid.xform transformation was used to transform our NMF coordinates into the reference brain atlas space, allowing us to assign brain region labels to our data and pool results across all fish.

### Multivariate linear regression analysis

We used multivariate linear regression to regress the activity of each neuron against 10 timeseries variables (six behavioral variables, three qREM state-related variables, and a constant offset term). Behavioral variables included speed, roll, pitch, left eye angle, right eye angle and turn bias (defined below). qREM state variables included a binary indicator function for qREM state, its integral, and its positive derivative (convolved with an exponential kernel, time constant = 1/15 * the duration of the qREM period). Collectively, these vectors comprise a 10-by-time matrix, *X,* such that we can define our model for a given neuron, *y_i_* as:

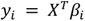

where *β_i_*is the 10-by-1 set of regression coefficients for this neuron which can be solved for as follows:

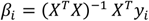

In this way, *β_i_* was fit for each neuron. We fit each cell using 10-fold nested cross validation. Briefly, data was divided into 10 non-overlapping time windows. Fits were performed on 90 % percent of the data and evaluated on the heldout 10 % percent. This was repeated 10 times to generate a prediction of the entire time series. Model performance was quantified using *R*^2^ between the predicted and actual activity in the held-out validation data. To test the significance of each specific regression variable, we fit additional models, using the same procedure, but in which regressor(s) of interest were temporally shuffled prior to model fitting. This procedure effectively broke the temporal relationship between the regressor and the neural activity, while preserving the number of degrees of freedom in the model. Thus, we could compare *R^2^* of the shuffled model fit directly to the full model. A regressor was deemed to contribute significantly to the variance of a neuron if, after shuffling, the reduction in mean *R^2^* across validation sets was larger than 2 times the standard error of *R^2^* across validation sets. We ran three such partial models: 1) left / right eye angles shuffled, 2) turn bias shuffled, 3) qREM variables shuffled. We used these models to identify the neurons encoding eye position, turn bias, and qREM, respectively.

### Definition of turn bias

We defined a behavioral regression variable based on animal heading in order to capture oscillatory turning dynamics. Turn bias was defined by convolving fish heading angle change with a Gaussian kernel with σ = 8 minutes. This resulted in a smoothed version of Δheading that remained stably positive or negative over the course of multiple minutes, reflecting sustained turn direction bias (Extended Data Fig. 5).

### In situ labeling of dbh positive cells

We performed imaging in freely moving fish as described above. We used chambers with confined areas (e.g., corners) as these were found to induce qREM more reliably.

We fixed and labeled fish with *in situ* hybridization^51,52^. Briefly, we first fixed the fish in 4 % PFA in PBST at 4°C overnight, then incubated for 2 to 10 min in methanol at −20°C, followed by rehydration with 50 % and 25 % methanol for 5 min each. We then incubated the fish overnight at 37°C with DNA probes designed to tile the dbh mRNA (Molecular Instruments, Inc). We then washed these probes and incubated overnight at room temperature with fluorescently labeled snap-cooled hairpins with an excitation wavelength of 514 nm (B2 amplifier, Molecular Instruments, Inc.). Finally, we removed the hairpins with 4 washes (2 x 5 min, 2 x 30 min and 1 x 5 min) in 5X SSCT. We mounted the fish in 2 % agarose and imaged the fluorescent labeling as a brain volume with a LSM 780, AxioObserver confocal microscope (Zeiss) with a LD LCI Plan-Apochromat 25x/0.8 Imm Korr DIC M27 objective at 1 μm × 1 μm × 2 μm resolution.

We used CMTK to align the confocal GCaMP stack with the averaged functional stack:

Step 1: cmtk make_initial_affine --centers-of-mass Functional.nii Confocal.nii initial.xform

Step 2: cmtk registration --initial initial.xform --dofs 6,12 –ncc or nmi --exploration 75 --accuracy 4 -o affine.xform Functional.nii Confocal.nii

Step 3: cmtk warp -o nonrigid.xform --nmi, smi or ncc --jacobian-weight 0 --accurate-e 18 --grid-spacing 80 -- energy-weight 1e-1 --refine 6 --coarsest 6 --ic-weight 0 --output-intermediate --accuracy 0.8 --initial affine.xform Functional.nii Confocal.nii

In cases for which the initial step didn’t give satisfying results, we preceded CMTK alignment with landmark registration using the “Name Landmarks and Register” plugin in FIJI. We applied the same transformation to the in situ staining data, and thresholded the resulting map to remove background and generate masks.

To extract single neuron traces from our functional data, we used a modified version of the NMF procedure described above. 2D NMF performed at 2 μm axial spacing leads to multiple subsampling of cells that span more than one axial plane. Thus, multiple axial ROIs can correspond to a single cell in the in-situ patterns. To resolve this, we instead separated the volume in sub-volumes of 31×56×56 pixels overlapping by 6 pixels in *x*, *y* and *z*. We first processed these volumes using PCA/ICA as described in^67^ which led to maps with well separated cells in 3D. We used these maps to obtain cell centroids (local maxima), and NMF to unmix the volume in the vicinity (6 μm) of these centroids as described above.

To compare the activity between wake and qREM, we averaged single neuron z-scored traces for each behavioral state, and then averaged each value per fish.

### Anatomical enrichment

To determine which brain regions were enriched for qREM encoding neurons, we computed the fold-enrichment of qREM neurons in each brain region relative to a random sample of neurons in the brain. That is, for a given brain region, *X*, we first determined the number of qREM positive neurons, *n_pos, region_*, and the number of total neurons imaged in the region, *n_total, region_*. These were used to define the proportion of qREM neurons in this brain region, *P(X)*.

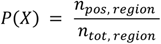

Next, we sampled *n_pos, region_* neurons from the pool of all recorded neurons in the experiment, across the entire imaging volume, and determined how many of these neurons were qREM positive, *n_pos, rand_*. We generated a null distribution, *P_0_(X)* by repeating this procedure 100 times.

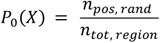

Finally, fold enrichment, *E(X),* was defined as the ratio of actual probability that a neuron in region *X* is qREM positive to the null probability of finding a qREM positive neuron in the brain.

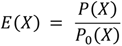

For a given fish, we defined regions as significant if *P(X) > P_0_(X)* in at least 95 % of random samples (p < 0.05). Across fish, a region was declared significant if at least 8/11 fish met this criterion. We ran this analysis for all brain regions available in the MapZebrain atlas^63^ as well as a custom defined region in the hindbrain based on the results in Fig. 5e.

### Exponential model fits to qREM activity

We used exponential model fits to quantify the dynamics of positively modulated qREM neurons within each qREM period. qREM states with durations < 3 min were excluded from this analysis. First, activity of each identified qREM neuron was linearly interpolated to remove missing values. Next, a qREM “trial”-by-time matrix was built by extracting the neuron’s activity during each qREM period and resampling along the time axis to 1000 data points, thus transforming activity into “relative” time within each qREM state. Each qREM state was then treated as a trial, and the mean across qREM “trials” was computed and normalized, and the resulting 1×1000 timepoint vector was fit with exponential models. We fit two exponential models to each cell, including an exponential decay model,

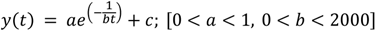

and a “1 minus exponential” model,

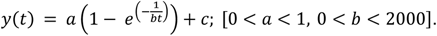

Models were fit by minimizing the squared error between true activity and predicted activity and performance was measured with *R^2^.* Individual neurons were classified as ramp up or ramp down according to which model performed better. No cross-validation was performed as our goal here was to categorize cells as ramp up or ramp down and to quantify their temporal dynamics.

### PCA of whole-brain neural activity

Neural activity was linearly interpolated and *z*-scored prior to performing PCA. In order to focus specifically on variance associated with qREM or wake, respectively, we performed two analyses. In the first analysis, the activity during each qREM period was extracted and time was downsampled such that each qREM period was described by a neuron x 10 time point matrix. We computed the mean of these matrices across qREM periods for each fish, performed PCA on this mean activity matrix, and projected the data from the full experiment onto the top two PCs of this space. In the second analysis we performed exactly the same procedure, but restricted to wake periods instead of qREM.

### Entanglement of state space trajectories

In order to measure how organized neural activity trajectories were in the PC space described above, we computed a measure of entanglement for each state trajectory. We defined entanglement as the number of times a given trajectory crossed over itself between the beginning and end of a given qREM state. Self crossings were defined by first approximating each qREM trajectory (first smoothed with Gaussian kernel, σ = 12.5 s) as a collection of piecewise linear functions for each consecutive sampling point (2 Hz sampling for neural data). That is, each trajectory was approximated by N-1 linear segments where N is the number of time points in the trajectory. For each linear segment, we determined how many times it intersected another segment from the collection. The total number of intersections was divided by 2 to count only unique crossings, and was then normalized by the duration of the given state trajectory. Measurements of qREM entanglement were performed in the qREM PCA space, and wake entanglement was measured in the wake PCA space.

### Relative time decoding from whole-brain activity

Population activity was used to decode relative time within each behavioral state. First, the activity of each neuron was linearly interpolated to remove missing values. Subsequently, *z*-scores were computed for the neural activity within each state, namely wake or qREM. We excluded short state periods which lasted less than 3 minutes. To decode the relative time within a specific state, we partitioned the *z*-scored neural activity and behavioral metrics for each state into 100 bins representing relative time within each state. To establish the ground truth for relative time, we defined a response variable that linearly increased from 0 to 1 within each state. To predict this response variable, we applied an elastic net linear model (sklearn package in Python)^68^ to the population neural activity. This model incorporated L1 and L2 penalty priors as regularizers. During the training phase of the decoder, data from all instances of a given behavioral state, except one held out instance, were utilized. Relative time decoding performance then was evaluated on the held-out state instance. After predicting relative time for all the periods, we employed a second stage of gain correction of relative time predictions. We fitted a common gain factor using the predictions of all except one held-out fish. We then gain-corrected the prediction of the held-out fish using the fitted gain factor. We repeated this for the predictions of all the fish.

An exhaustive search per fish was conducted to identify the optimal hyperparameters for both qREM and wake states, as illustrated in Extended Data Figure 7a-b. To evaluate the decoding performance against chance level models, we temporally shuffled the neural activity within a given state while preserving local temporal correlation. To determine whether a neuron consistently contributed to the decoding of relative time, we selected neurons whose median decoding weights across model fits were greater than 0 or less than 0. Because of the L1 component of model regularization, many neurons received a weight of exactly 0 during model fitting. Subsequently, we defined ramp up neurons as those having consistent positive decoding weights (median coefficients > 0) and ramp down neurons as those having consistent negative decoding weights (median coefficients < 0) across different training sets.

### Statistical analysis

Sample sizes, statistics, and statistical tests are reported in the figure legends and main body of the results. For comparisons of behavior and neural activity across different experimental or internal state conditions, we used Wilcoxon rank sum and Wilcoxon signed rank tests. In all cases, two-sided tests were performed. In all cases where multiple observations (e.g., neurons) were collected from one animal, we first computed the mean statistic within animal before performing the statistical test across conditions. While this approach slightly reduces statistical power, it leads to more conservative estimates of p-values. Sample size was not predetermined by a statistical method. The experiments were not randomized and were not performed with blinding.

### Animal care and transgenic lines

Experiments were performed in accordance with the Animal Welfare Office at University of Tübingen and the Regierungsprasidium. The majority of the imaging experiments used Tg (elavl3:H2B-GCaMP6s+/+ or elavl3: GCaMP6s+/+) with *nacre* (*mitfa*−/−) at 5-8 dpf. In a subset of imaging experiments, we used Tg (elavl3:H2B-GCaMP8s+/+) with *nacre* (*mitfa* −/−) 5-8 dpf. Behavioral experiments used *nacre* (*mitfa* −/−) at 5-8 dpf. For one set of behavioral experiments, we used Tg (*lakrtiz* −/−) at 5-6 dpf. Fish were reared on a 14/10 hr light/dark cycle at 25°C. They were maintained in group of 20 and fed dry food daily.

### Data availability

All data that contributed to the findings of this work will be made available following publication upon reasonable request.

### Code availability

All analysis software was written in Julia 1.7-1.8, Python 3.10, and CUDA C using the Julia package ecosystem (Optim.jl, HDF5.jl, JLD2.jl, PyPlot.jl, CUDA.jl, Images.jl) and Python package ecosystem (scikit-learn, scipy, pandas, matplotlib, numpy). GPU acceleration was implemented for fish tracking, as well as online and offline image registration.

## ACKNOWLEDGEMENTS

We wish to thank Dr. Herwig Baier for providing us with the *Tg(lakritz−/−)* fish line and Dr. Aristides Arrenberg and Dr. Emre Aksay for valuable discussion about eye movements in larval zebrafish. We are thankful to our lab members for their feedback on the manuscript and the machine and electronic workshops at the Max Planck Institute for Biological Cybernetics, Tübingen, Germany for helping us with the experimental setup.

## AUTHOR CONTRIBUTIONS

JML, DNR, and VC conceived the project. VC and CH collected and analyzed the primary behavioral and neural data in this manuscript, CH created cluster computing pipelines for data analysis, SA collected and analyzed the lakritz and HCR experiments, and LSS contributed preliminary data on qREM and analysis of brain activity during flow. JML and DNR provided guidance on data collection and analysis. DNR designed the hardware and software for imaging systems and primary data processing pipelines used in the manuscript, and all authors contributed to their testing and implementation. All authors contributed to the writing of the manuscript.

## MATERIALS AND CORRESPONDANCE

All correspondence should be addressed to the corresponding authors

## EXTENDED DATA FIGURES

**Extended Data Figure 1.**
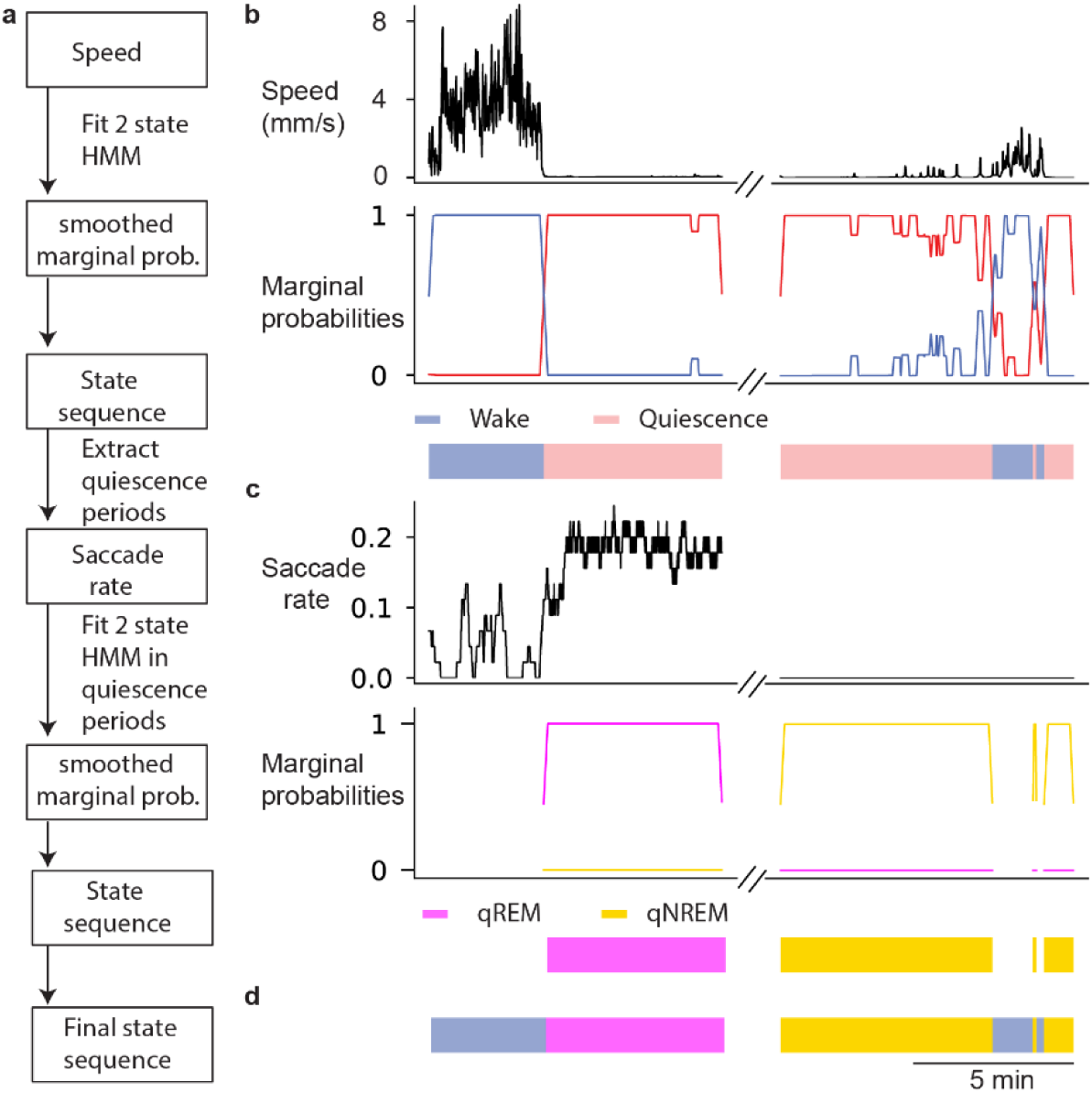
Hierarchical HMM workflow for behavioral state labeling. **a,** Pipeline for two stage HMM fitting. The first stage assigns labels to wake and quiescence periods based on speed and the second stage assigns labels to substates of quiescence, qREM and qNREM, using saccade rate. **b**, Illustration of stage 1 of the pipeline. Speed trace on top, marginal probabilities (posteriors) of wake and quiescence states in the middle and state sequence on the bottom. **c**, Illustration of stage 2 of the pipeline. HMM fitting is only performed on quiescent periods identified in stage 1. Saccade rate trace on top, marginal probabilities (posteriors) of qREM and qNREM substates of quiescence in the middle and state sequence on the bottom. **d**, Final state sequence after both stages of HMM fitting.

**Extended Data Figure 2.**
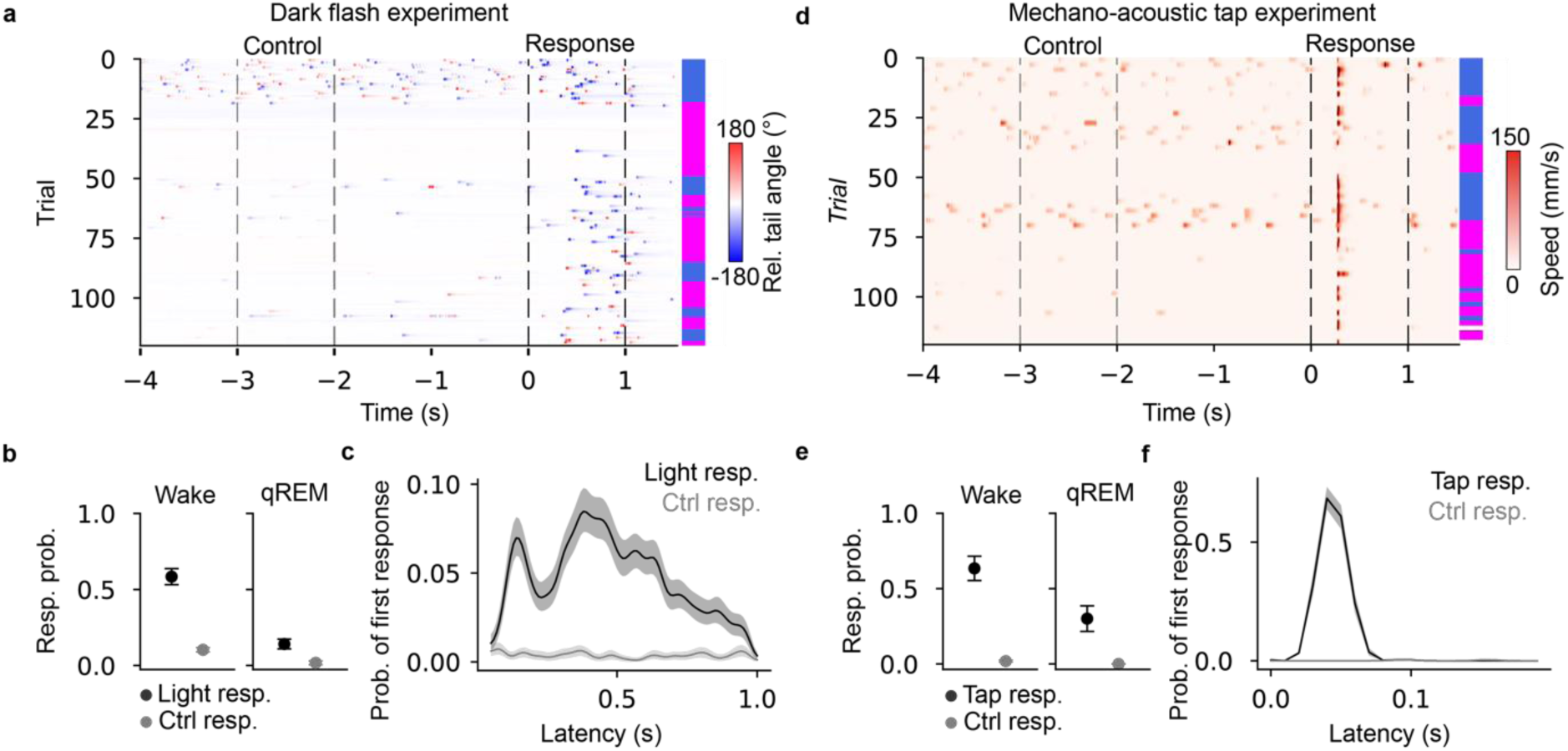
Overall movement level does not explain state-dependent arousal threshold. **a,** Raster plot of relative tail angle in one example fish. Trials are sorted according to the time in which they occurred in the experiment. Behavioral state on each trial (qREM vs. wake) is indicated by the vertical hypnogram (wake: blue, qREM: magenta) on the right. “Control” response window highlights a randomly chosen 1 s window used to perform a matched control analysis of the true response window, defined as the light off period. **b,** Response probability as a function of behavior state (wake vs. qREM) and analysis window (control vs. actual response window). In both behavioral states, response probability was significantly higher during the actual response window (wake: p = 3.0e-7, qREM: p = 0.001, n = 24 fish, Wilcoxon signed rank test), indicating that wake responses are not simply a consequence of higher baseline locomotion. **c,** Distribution of response latencies detected during the actual (black) and control (grey) analysis windows. Shading represents bootstrapped 95 % confidence interval calculated in a 50 ms sliding window. **d,** Raster plot of speed in one example fish. Trials are sorted according to the time in which they occurred in the experiment. Behavioral state on each trial (qREM vs. wake) is indicated by the vertical hypnogram on the right. “Control” response window highlights a randomly chosen 1 s window used to perform a matched control analysis of the true response window, defined as one second following the solenoid tap. **e,** Response probability as a function of behavior state (wake vs. qREM) and analysis window (control vs. actual response window). In both behavioral states, response probability was significantly higher during the actual response window (wake: *p* = 0.01, qREM: *p* = 0.01, n = 8 fish, Wilcoxon signed rank test), indicating that wake responses are not simply a consequence of higher baseline locomotion. **f,** Distribution of response latencies for responses detected during the actual (black) and control (grey) analysis windows. Shading represents bootstrapped 95 % confidence interval calculated in a 10 ms sliding window.

**Extended Data Figure 3.**
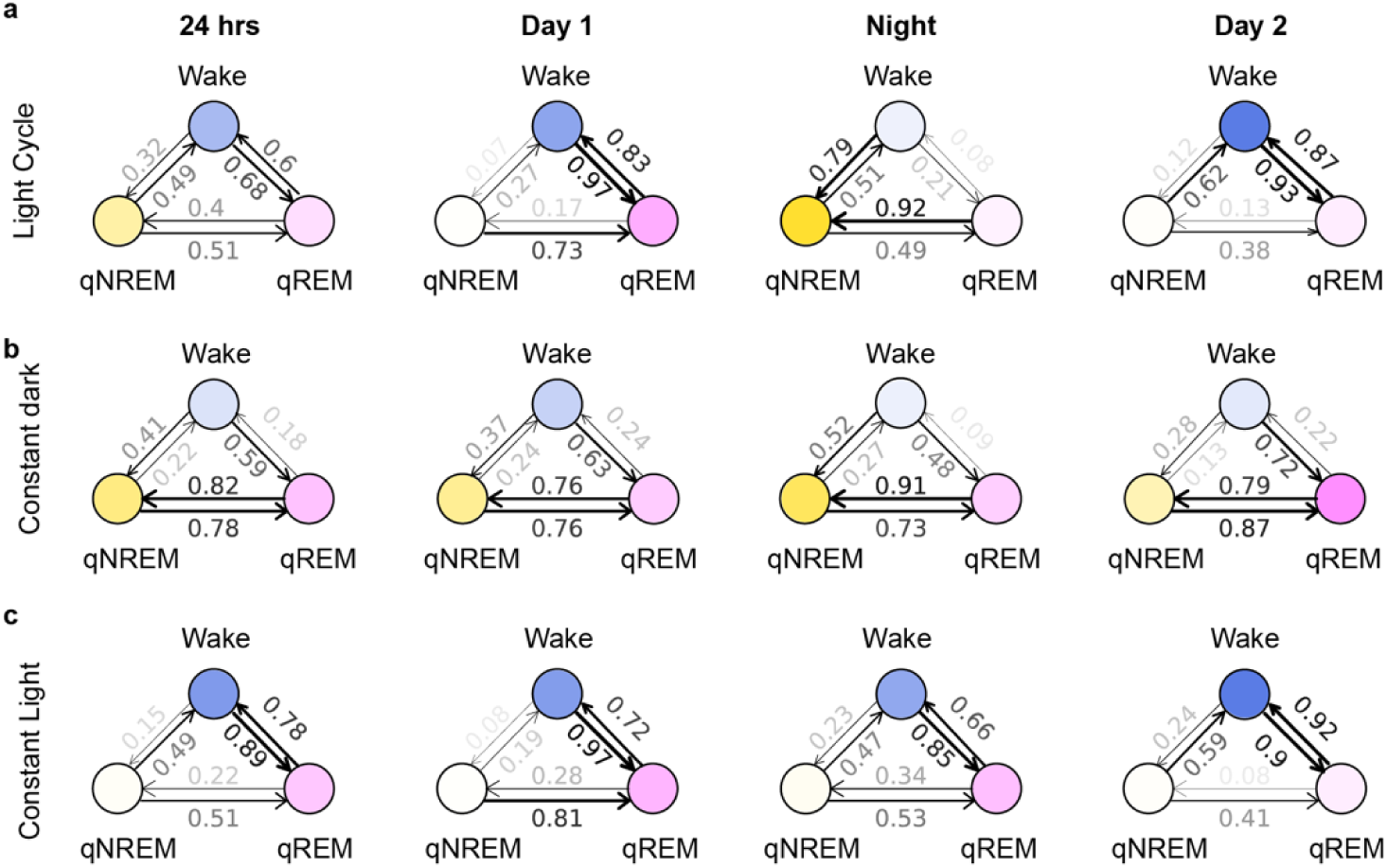
Transition probabilities between wake, qREM and qNREM states. Transition probabilities between wake, qREM and qNREM for light cycled (**a**, n = 53 fish, 14 hr light / 10 hr dark), constant dark (**b**, n = 27 fish), and constant light (**c**, n = 51 fish) conditions over 24 hr. Arrows indicate transition direction, and their widths are proportional to their respective transition probabilities. Transparency of each circle is proportional to the time occupied by the respective state. Transition probabilities in the left column are computed from the whole 24hr experiments with the exclusion of states shorter than 1 minute. These 24hr experiments are broken into day 1, night, and day 2 according to the entrained circadian cycle during rearing conditions (14 hr light / 10 hr dark). Transition probabilities for these periods are in the middle and right columns.

**Extended Data Figure 4.**
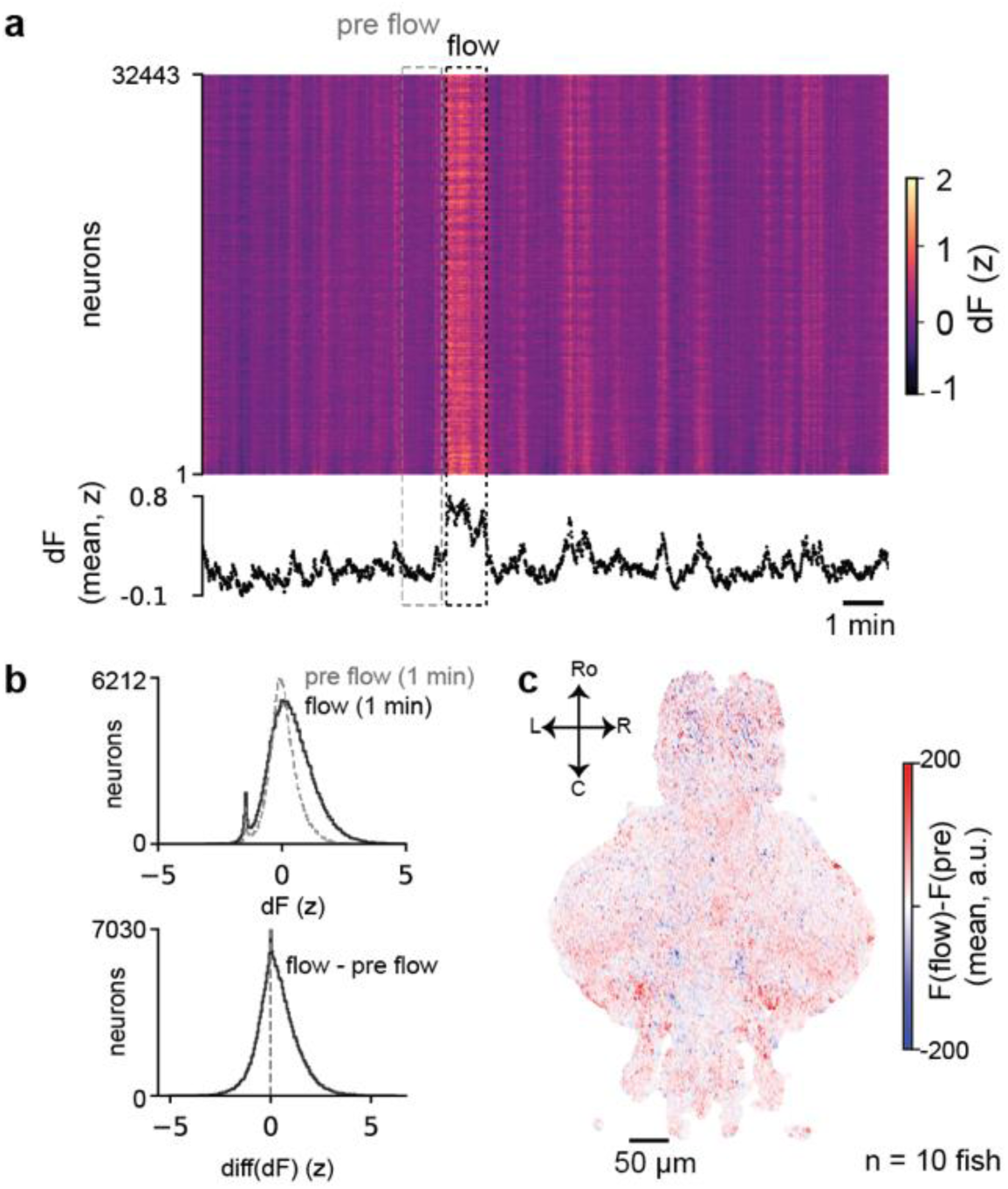
Flow triggers transient increase in brain activity. **a,** Raster plot showing brain-wide neural activity of a representative example fish experiencing 1 min of flow (dashed black box). Below: mean activity across all shown neurons. **b**, Histogram of neural activity (z-scored, n = 10 fish) over a period of 1 min before flow (grey dashed) and 1 min during flow (black). Below: histogram of difference in activity between pre and during flow for each neuron (mean of 1 min during flow “minus” mean of 1 min before flow) **c,** Mean difference in fluorescence of flow and pre flow period for each neuron across fish (266,326 neurons, n = 10 co-registered fish, dorsoventral mean projection).

**Extended Data Figure 5.**
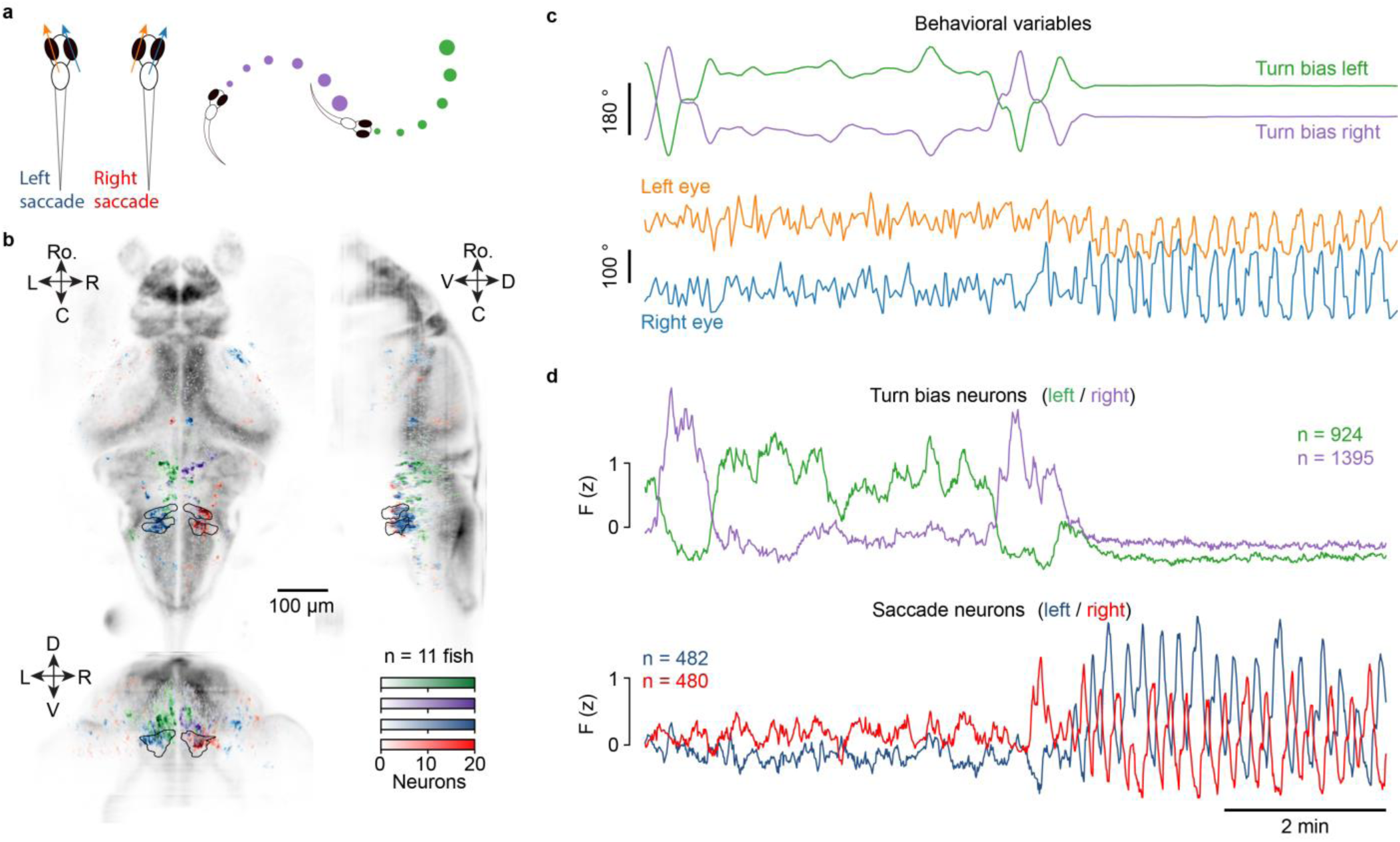
Distinct neural populations control eye saccades and turn bias. **a,** Eye angles and turn bias were measured from behavioral tracking. Left and right eye angles (orange / blue arrows) are defined as the angle between the eye’s major axis and the animal’s heading. An example of leftward (blue) and rightward saccades (right) are shown. Left (green) and right (purple) turn bias, illustrated as the size of the dots, are defined as the Gaussian smoothed (σ = 8 min) heading change over time. Thus, continuous turning in the leftward direction leads to larger values of left turn bias and vice versa for continuous right turning. **b,** Density map of all categorized neurons across n = 11 fish registered to the mapZebrain atlas (**Methods**). Colors correspond to the behavioral metrics illustrated in **a.** Black tracing indicates the contour of the abducens nucleus. **c,** 15 min example traces of behavioral variables (eye angle, and turn bias) that were input to the regression model. **d,** Mean activity traces of the neural populations corresponding to each behavioral variable. Each trace is a mean over all neurons belonging to each category, identified by linear regression (**Methods**).

**Extended Data Figure 6.**
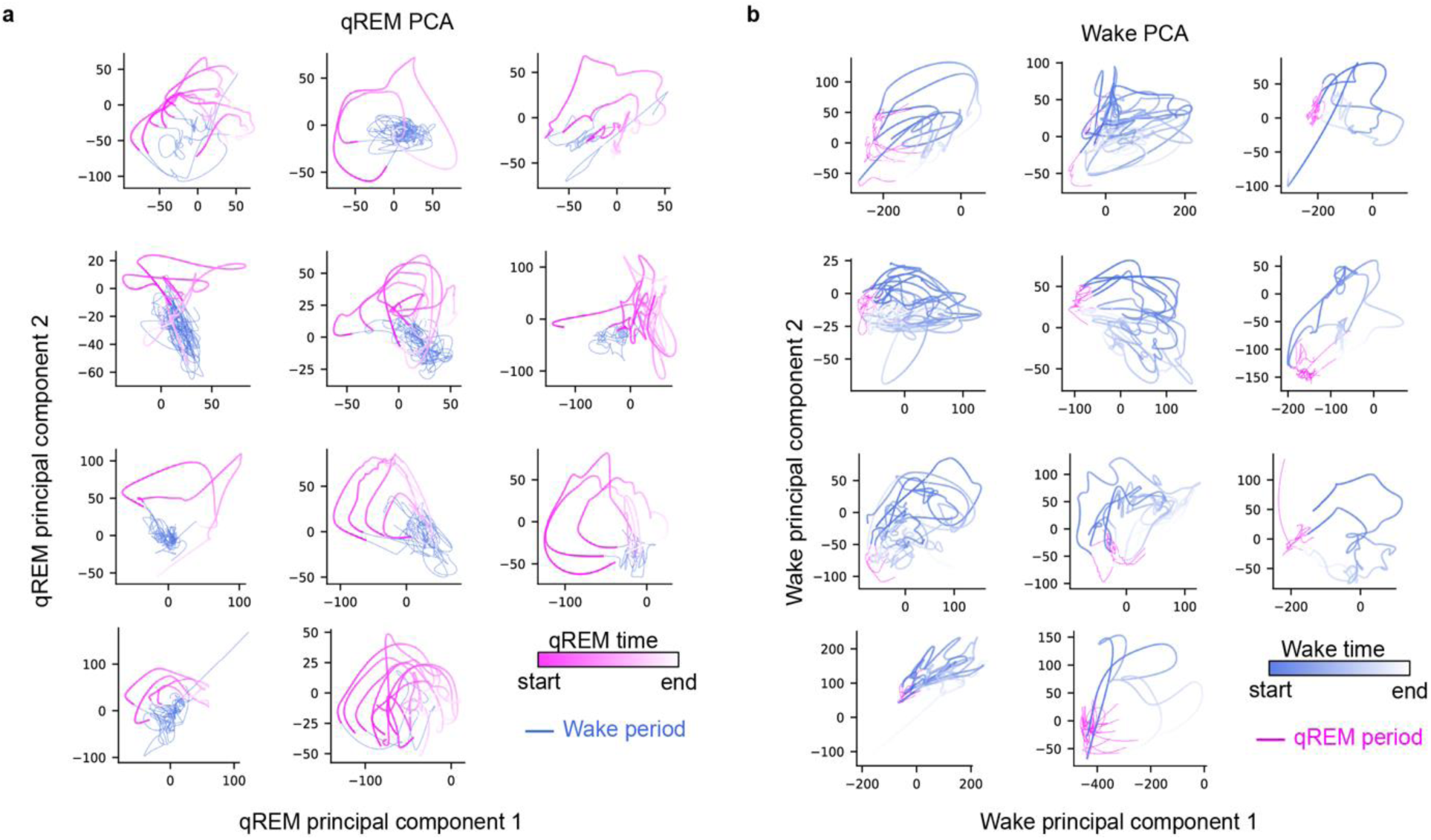
Unsupervised discovery of smoothly evolving state trajectories. Whole-brain PCA was performed only during qREM periods (**a**) or wake periods (**b**). All timepoints are projected onto the top two PCs of the respective space **(Methods).** In the qREM space (**a**), time within each qREM period is represented by the colormap (magenta to white). All wake time points are colored blue. In the wake space (**b**), time within each wake period is represented by the colormap (blue to white). All qREM time points are colored magenta.

**Extended Data Figure 7.**
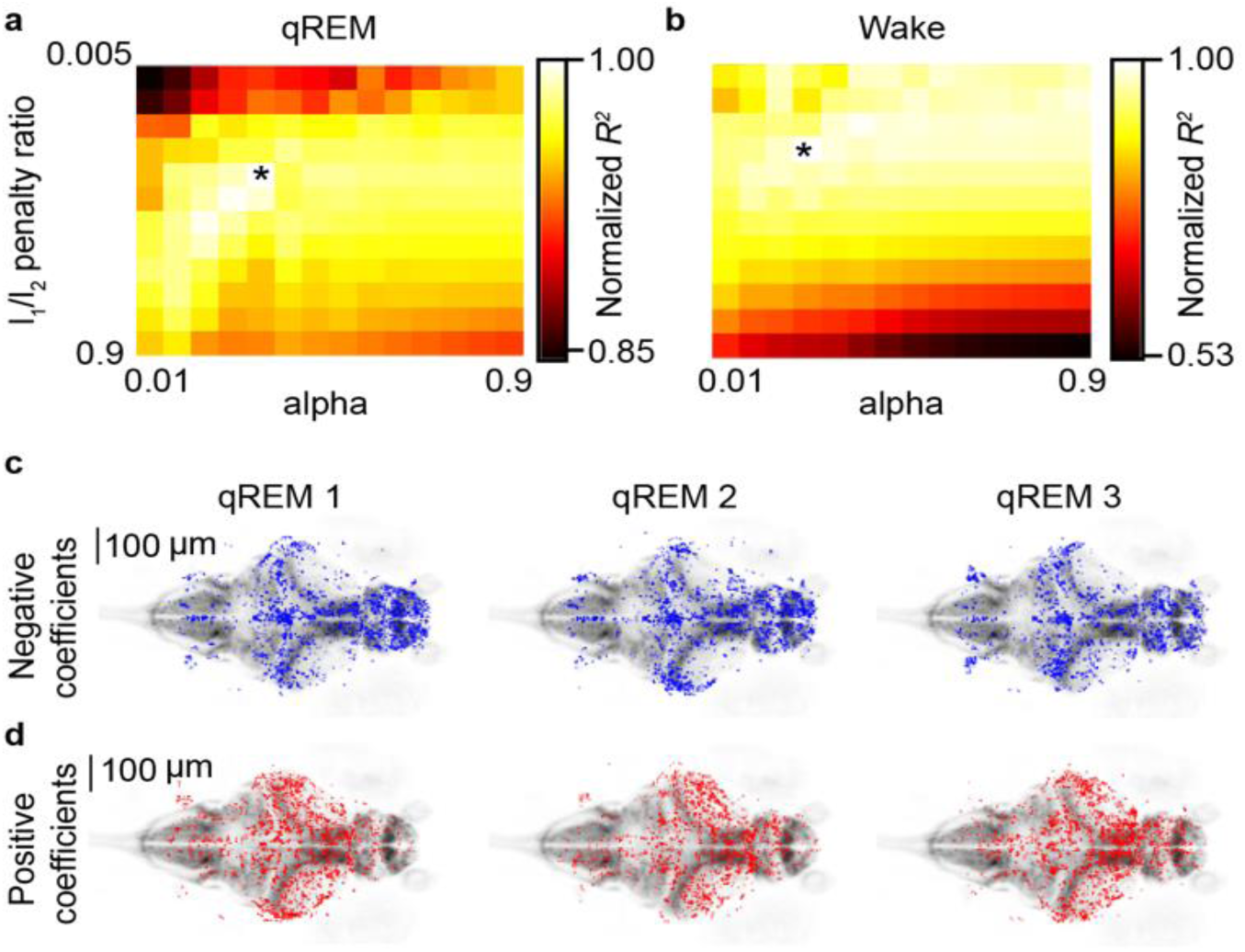
Hyperparameter optimization and neuron selection. **a-b**, Normalized coefficient of determination (*R^2^* score, n = 11 fish) heatmap for *L_1_/L_2_* penalty ratio and penalty scaling value (alpha) for decoding relative time in qREM (**a**) or wake (**b**) periods. Asterisk (*) indicates the maximum *R^2^* score (*l_1_/l_2_* penalty ratio = 0.2, alpha = 0.05) for qREM (**a**) and maximum *R^2^* score (*l_1_/l_2_* penalty ratio = 0.1, alpha = 0.04) for wake (**b**). **c-d**, An example dataset. Neurons with negative (**c**) or positive (**d**) decoding weights in each qREM training set. For example, qREM 1 training set is qREM 2 and qREM 3.

**Extended Data Figure 8.**
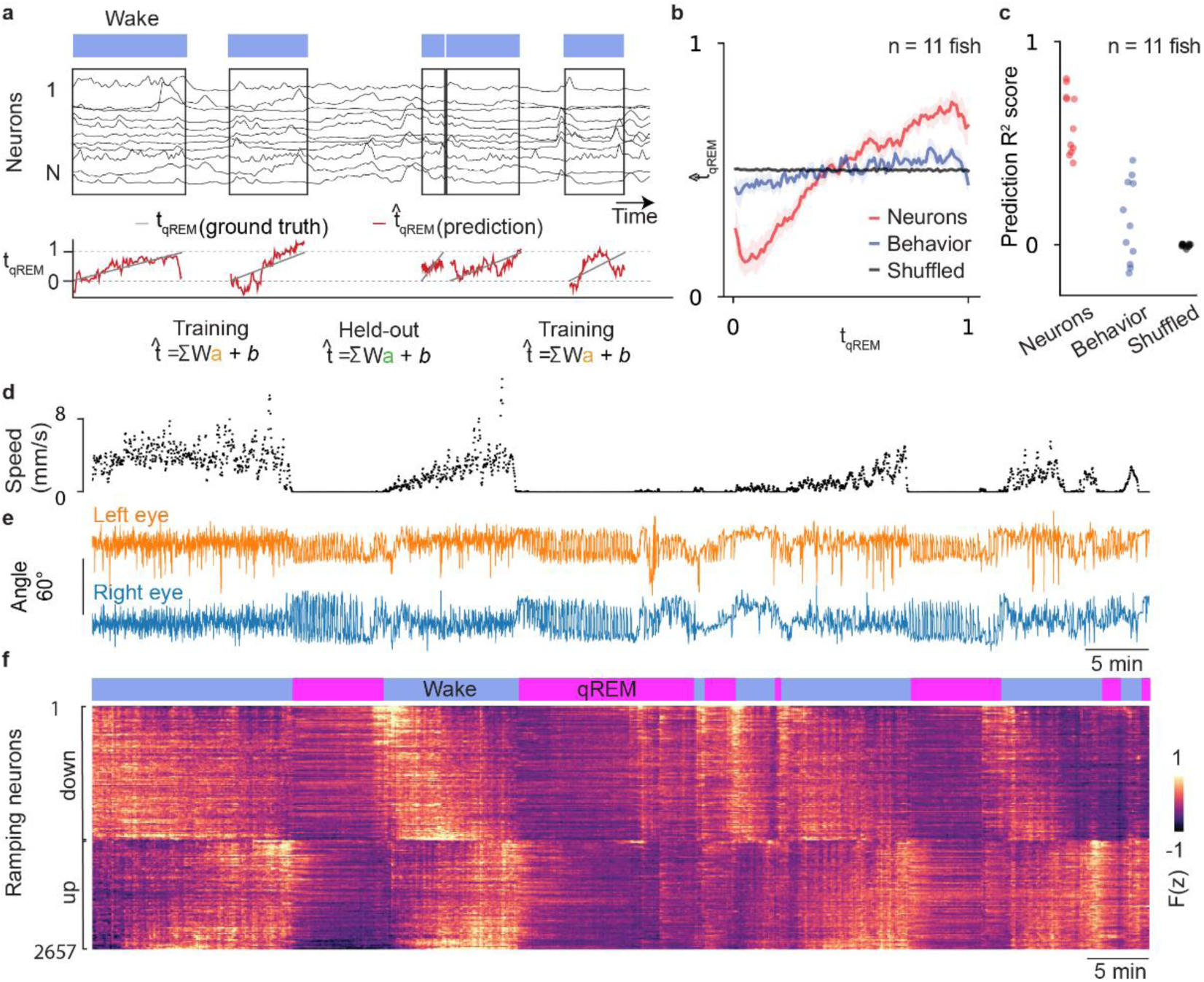
Relative time decoding during wake periods. **a**, Relative time decoding pipeline. Top: wake state labels and activity of 11 example neurons smoothed with a Gaussian kernel (σ = 25 s). The linear decoder was trained on all wake states except for one. Trained decoder weights were then applied to the held-out wake period to predict relative time. This procedure was repeated for each wake state in order to generate predictions for the entire dataset. Bottom: Relative time prediction (red) in each wake state compared with true relative time (grey). **b**. Relative time prediction (mean ± s.e) using only neural activity (red), behavioral variables (blue), or shuffled neural activity (black) (n = 11 fish). **c**. Decoder performance was quantified by measuring *R^2^* between true relative time and predicted related time (neural *R^2^* = 0.60 ± 0.05, shuffled *R^2^* = −0.01 ± 0.01, behavior *R^2^* = 0.12 ± 0.06, n = 11 fish). **d-e** Speed (mm/s) (**d**), left eye (**e**, orange), and right eye (**e**, blue) angle of an example freely swimming larva. **f**. Top: Behavioral state labels (blue: wake, magenta: qREM) throughout the experiment. Bottom: Raster map of z-scored neural activity for neurons in the example animal that significantly contributed to relative time decoding during wake state: ramp-down neurons (1470, top rows), which had negative decoding weights, and ramp-up neurons (1187, bottom rows), which had positive decoding weights.

## SUPPLEMENTARY INFORMATION

**Supplementary Video 1 | Example wake, qREM, and qNREM behavior.** Behavior of one fish during the wake state (left), qREM state (middle), or qNREM state (right). Each panel highlights 45 s of behavior from each state. Insets of the animal’s eyes are at 6X zoom relative to the overall behavior chamber. Below, the tracked eye angles are shown for each example data excerpt.

**Supplementary Video 2 | Brain-wide activity at the transition between wake and qREM.** Raw fluorescence data, normalized to mean wake activity in the 2 min preceding qREM onset, is shown for one example fish at the transition between wake and qREM. Normalized activity at each time point is averaged across all wake to qREM transitions (n = 3 transitions in this fish). Each panel represents a different slice through the dorso-ventral axis (reading left to right and top to bottom: dorsal to ventral).

